# 40S ribosomal subunits scan mRNA for the start codon by one-dimensional diffusion

**DOI:** 10.1101/2024.12.30.630811

**Authors:** Hironao Wakabayashi, Mingyi Zhu, Elizabeth J. Grayhack, David H. Mathews, Dmitri N. Ermolenko

## Abstract

During eukaryotic translation initiation, the small (40S) ribosomal subunit is recruited to the 5′ cap and subsequently scans the 5′ untranslated region (5′ UTR) of mRNA in search of the start codon. The molecular mechanism of mRNA scanning remains unclear. Here, using GFP reporters in *Saccharomyces cerevisiae* cells, we show that order-of-magnitude variations in the lengths of unstructured 5′ UTRs have a modest effect on protein synthesis. These observations indicate that mRNA scanning is not rate limiting in yeast cells. Conversely, the presence of secondary structures in the 5′ UTR strongly inhibits translation. Loss-of-function mutations in translational RNA helicases eIF4A and Ded1, as well as mutations in other initiation factors implicated in mRNA scanning, namely eIF4G, eIF4B, eIF3g and eIF3i, produced a similar decrease in translation of GFP reporters with short and long unstructured 5′ UTRs. As expected, mutations in Ded1, eIF4B and eIF3i severely diminished translation of the reporters with structured 5′ UTRs. Evidently, while RNA helicases eIF4A and Ded1 facilitate 40S recruitment and secondary structure unwinding, they are not rate-limiting for the 40S movement along the 5′ UTR. Hence, our data indicate that, instead of helicase-driven translocation, one-dimensional diffusion predominately drives mRNA scanning by the 40S subunits in yeast cells.

## Introduction

In eukaryotes, translation typically begins at the AUG codon that is closest to the 5′ end of mRNA (1). During eukaryotic translation initiation, initiation factor <u>eIF4E</u> (<u>e</u>ukaryotic Initiation <u>F</u>actor 4E) recognizes the m^7^Gppp cap structure at the 5′ end of the mRNA and recruits two other initiation factors, eIF4G and eIF4A, by forming a complex called eIF4F. eIF4F recruits the small (40S) ribosomal subunit preassembled with initiator tRNA and initiation factors 1, 1A, 2, 3 and 5 (2). After recruitment to the 5′ end of mRNA, the small ribosomal subunit is believed to move along the 5′ UTR (3–6) in search of the start codon.

mRNAs vary in their translational efficiencies by at least two orders of magnitude (7–10). The 5′ UTR is thought to play a critical role in regulating translation initiation through variety of mechanisms, many of which are not well understood. For instance, 5′ UTR length and secondary structure may regulate translation by changing the rate of mRNA scanning by the 40S. Indeed, introduction of stable stem-loops in the 5′ UTRs was shown to reduce translation presumably by hampering 40S movement from the cap to the start codon (11–15). Mutations affecting secondary structure in the 5′UTR of BRCA1 mRNA were implicated in breast cancer development, linking 40S scanning and human disease (16). mRNA scanning by the 40S and the 5′ UTR structure were also associated with malignant transformation caused by upregulation of eIF4F (17). Although eIF4F recognizes the cap structure, which is common for most mRNA, upregulation of eIF4F enhances protein synthesis from just a subset of mRNAs (18). Many of these mRNAs were thought to have extensive secondary structure in the 5′ UTR, which could hinder 40S scanning (18,19).

Although 40S scanning is evidently involved in translational control, the mechanism by which the 40S subunit moves along the 5′ UTR and the regulation of 40S scanning by 5′ UTR remain obscure. It is commonly assumed that scanning is driven by one of the core initiation factors, DEAD-box RNA helicases eIF4A (20,21). Other translational helicases, such as Ded1 (DDX3 in mammals), are thought to contribute to 40S scanning through structured 5′ UTRs (22,23).

Helicase-driven models of 40S scanning tend to conflate secondary structure unwinding and 40S movement along the 5′ UTR that might be mechanistically distinct processes. 40S migration along the 5′ UTR may also occur by factor-independent diffusion (24–26).

Specifically, entanglement of secondary structure unwinding and 40S movement along the 5′ UTR hampers understanding regulation of 40S scanning by 5′ UTR length. When compared to ORFs and 3′ UTRs, the length of 5′ UTRs is relatively short in diverse taxonomic classes of eukaryotes (27,28). For example, median 5′ UTR lengths in yeast and human cells are ∼50 and 200 nucleotides, respectively (27). Although 5′ UTRs tend to be relatively short, experiments with translational reporters vary in the degree to which translation is reduced by lengthening the 5′ UTR (28–35). The discrepancy between different studies might stem from superimposition of the effects of 5′UTR length and secondary structure.

To overcome this problem, we employ intrinsically unstructured sequences and re-examine contributions of the 5′ UTR length and secondary structure to translation initiation in yeast cells. We find that extending the 5′ UTR of the GFP reporter tenfold over the median yeast 5′ UTR length moderately reduces translational efficiency. We also find that loss-of-function mutations or deletions of translational helicases eIF4A, Ded1 and Slh1 similarly affect reporter mRNAs with short and long 5′ UTRs, indicating that these helicases are not rate-limiting for 40S movement along 5′ UTRs. Based on these observations, we hypothesize that, at least in yeast, 40S scanning might be predominantly driven by one-dimensional diffusion.

## Materials and Methods

### Reagents

Oligonucleotides were ordered from IDT. Enzymes were from NEB. Other chemicals and standard reagents were ordered from Fisher Scientific.

### Biological resources

Yeast *Saccharomyces cerevisiae* strains used in this work are listed in Table 1.

**Table 1.**
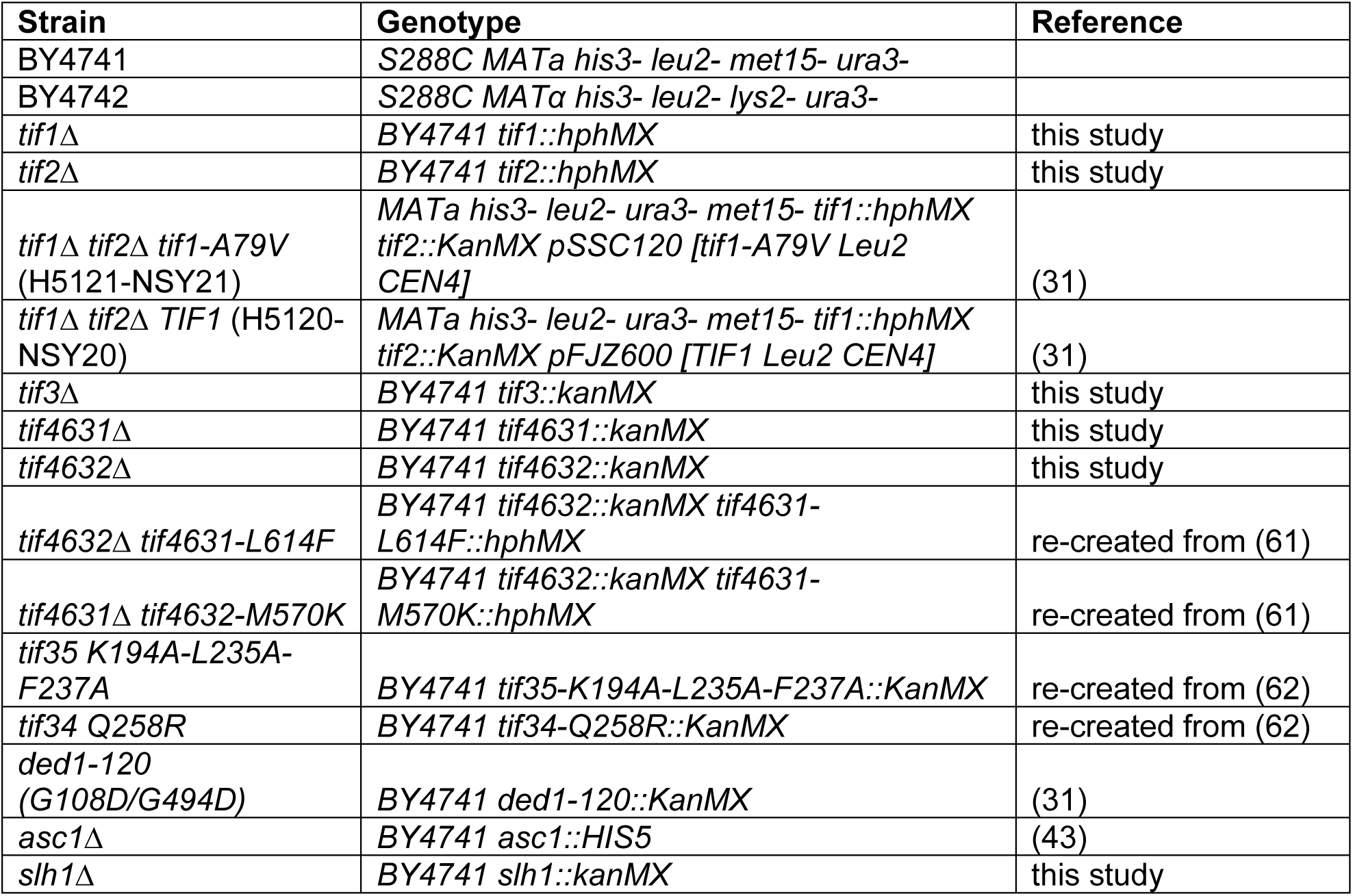
Yeast strains used in this work.

### RNA-ID constructs design and preparation

A GFP reporter containing UTRs from yeast RPL41B gene was synthesized by GenScript. The GFP ORF contains an N-terminal 3C site-HA epitope-His6-encoding sequence upstream of superfolder GFP sequence. Non-repetitive unstructured sequences (NUS) sequences were designed using the orega tool (optimizing RNA ends with a genetic algorithm) of the RNAstructure software package (https://rna.urmc.rochester.edu/RNAstructure.html) as previously described (36). Orega evolves sequences using a scoring function that penalizes basepair formation and encourages a diversity of sequence motifs as quantified by linguistic complexity(37,38). Default parameters were used for orega. By default, 1000 iterations of evolution occur, but calculations were restarted for additional sequence evolution when sequences were still predicted to have basepairs. Basepair probabilities were estimated using the partition function in RNAstructure (39).

We used the Gibson assembly method combined with conventional restriction digestion and ligation to construct RNA-ID DNAs with various sequences (^50^NUS ∼ ^500^NUS, ^300^NUS2, ^300^NUS + SL, ^50^NUS3 + ^122^Struct, ^50^NUS4 + ^122^Struct) at the 5’ end of the sfGFP-ORF sequence. Primers and ultramer oligo nucleotides, whose sequences can be found in Supplementary Tables 2-14, were synthesized at Integrated DNA Technologies (IDT, Coralville, IA). Detailed procedures including the production of ^122^Struct was described in Supplementary Methods.

### Yeast transformation with RNA-ID reporters

BY4741, BY4742, or mutant yeast strains were transformed with RNA-IDs constructs as previously (40). To obtain a linear DNA fragment containing the RNA-ID reporter flanked on each end by sequences homologous to *ADE2* locus, the RNA-ID plasmid was digested StuI restrictase (NEB) and then subjected to agarose-gel purification.

To prepare competent yeast cells for transformation, yeast cells were growth in YPD liquid media to OD_600_ between 1.3 and 2.5 (log phase). For each transformation, 10 mL of cell culture was pelleted by centrifugation at 3000 rpm for 5 min. Pellets were washed twice with 1 ml of 100 mM LiOAc. Next, cells were resuspended in 110 μl of 100 mM LiOAc containing salmon sperm DNA at concentration of 1 mg/ml, and then incubated with 300 ng of linearized DNA at 30°C for 15 minutes. Next, 600 μl of a LiOAc-PEG solution (8 volumes of 60% PEG 3350 to 1 volume of 1 M LiOAc to 1 volume of ddH_2_O) were added. After incubation at 30°C for 30 minutes, cells were mixed with 68 μl of DMSO and incubated at 42°C in for 15 minutes. After transformation, cells were pelleted at 1500 rpm for 7 minutes, resuspended in 1 ml of YPD media and incubated at 30°C for 1 hour while shaking. After this recovery period, 250 μl of cells were growth on a SC-met^‒^ or SC-ura^‒^ plate at 30°C for 2 days. Individual colonies were streaked onto another SC-met^‒^ or SC-ura^‒^ plate and growth at 30°C for 2 days.

### Flow cytometry

Cells from a single colony were inoculated into 5 ml of YP media supplemented with 2% raffinose, 2% galactose and 80 mg/L adenine and grown overnight at 30°C. An aliquot of overnight culture was diluted to final volume of 5 ml with the same media to obtain OD_600_ of 0.2 and then grown for 4–6 h at 30°C to OD_600_ of ∼0.8 (Dean and Grayhack, 2012). Temperature sensitive mutants were grown in YP media containing 2% raffinose and 80 mg/L adenine (i.e. without galactose) overnight at 30°C. An aliquot of overnight culture was then diluted to final volume of 5 ml with the same media supplemented with 2% galactose to obtain OD_600_ of 0.2 and then grown for 4-18 h at the non-permissive temperature (either 18 or 37 °C) to OD_600_ of ∼0.8. 500 μl of resulting cell culture was transferred to a 5-mL polystyrene round-bottom tube (Falcon) and kept on ice for an hour. Flow cytometry was performed for three independent isolates of each strain using LSR Fortessa flow cytometer (BD Biosciences, Franklin Lakes, NJ) (Dean and Grayhack, 2012). A reference strain carrying a modified RNA-ID reporter with only a 6 nt-long 5’ UTR (GO control) is used as the primary control in all flow cytometry experiments. For RNA-ID GO control, both GFP and RFP channels were adjusted to 26,000 AU. Flow cytometry data were analyzed using Cytobank flow cytometry data analysis platform (41).

### Yeast gene deletion clone generation

We created deletion strains in a BY4741 background. To create *tif1*Δ and *tif2*Δ, yeast cells were transformed with a DNA amplicon containing KanMX DNA, which encodes bacterial aminoglycoside phosphotransferase for kanamycin resistance, flanked by 500 nt-long sequences homologous to the sequences flanking either *tif1* or *tif2* ORF in yeast genome. To obtain *tif3*Δ, *tif4631*Δ, *tif4632*Δ and *slh1*Δ, yeast cells were transformed with a KanMX-containing cassette amplified from respective deletion strain from the single-gene deletion collection (42) provided by Eric Phizicky (University of Rochester). Primer sets used for each amplicon are listed in Supplementary Table 3. Amplicons were agarose-gel-purified and used for transformation to BY4741 cells by the method descried above. In order to obtain the *asc1*Δ strain, we used the YJYW17 strain (BY4741, asc1::HIS5 + pASC1-*URA3*), in which the *asc1* ORF is replaced with the HIS5 cassette and complemented with Asc1-expressing *URA3* plasmid (43). This strain was cultured on 5-FOA-URA-SD plate a 30°C for 2 days to eliminate Asc1-expressing *URA3* plasmid.

### Yeast gene mutation clone generation

We produced yeast mutant clones, which contain mutations *tif4631-L614F*, *tif4632-M570K*, *ded1-120*, *tif34-Q258R*, or *tif35-K194A-L235A-F237A* according to the method described elsewhere (44). BY4741 cells were transformed with two DNA cassettes. One cassette contained the ORF of interest with respective amino acid substitutions flanked by ∼500 nt and ∼300 nt-long sequences homologous to the sequences flanking the 5′ and 3′ ends of the ORF in yeast genome. The 3′ flanking sequence was followed by ∼500 nt of KanMX or HgrB DNA. The second cassette contained full-length KanMX or HgrB DNA followed by 500 nt-long sequence homologous to the respective sequence of yeast genome positioned 300 nt downstream of the ORF of interest. These cassettes were excised by restriction enzymes from the plasmids listed in Supplemental Methods. Mutations were verified by sequencing of respective ORFs in genomic DNA.

### Other procedures

Yeast genomic DNA (gDNA) purification, RT-qPCR for GFP and RFP mRNAs, SDS-PAGE analysis with fluorescence detection, 5′RACE analysis and examining yeast growth via spot assay were done using standard procedures described in **Supplementary Materials**.

## Results

### Simplified models describing dependence of protein synthesis on the length of the 5′ UTR

To analyze putative experimental outcomes of examining dependence of protein synthesis on 5′ UTR length using GFP reporters, we considered a simple mathematical approximation: the amount of translated protein is inversely proportional to the time needed to synthesize one GFP molecule. The time needed to synthesize one GFP molecule accounts for the duration of 40S scanning, which depends on the length of the 5′ UTR (*L*), and sum of all other steps of initiation, elongation and termination, which do not depend on the 5′ UTR length (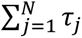 in equations 1-2). In this simplified approximation, mRNA half-lives and GFP protein degradation rates are assumed to be constant, independent of 5′ UTR length and unaffected by the overall rate of GFP synthesis. The duration of 40S scanning should scale linearly with 5′ UTR length if scanning is driven by a translocase/helicase (29):

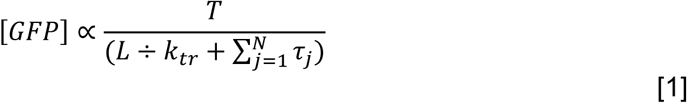

where *T* = total time, *L* = length of the 5′ UTR, *k_tr_* = rate of 40S translation along the 5′ UTR, and *τ_j_* = duration of *N* translation steps other than 40S scanning that are independent of 5′ UTR length.

Alternatively, scanning time should scale as the square of 5′ UTR length if scanning occurs by diffusion (29):

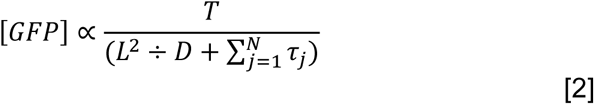

where *D* = 40S diffusion coefficient.

The reciprocal relationship between the amount of synthesized protein and 5′ UTR length is expected for both translocation and diffusional mechanism of scanning (Fig. 1).

**Figure 1.**
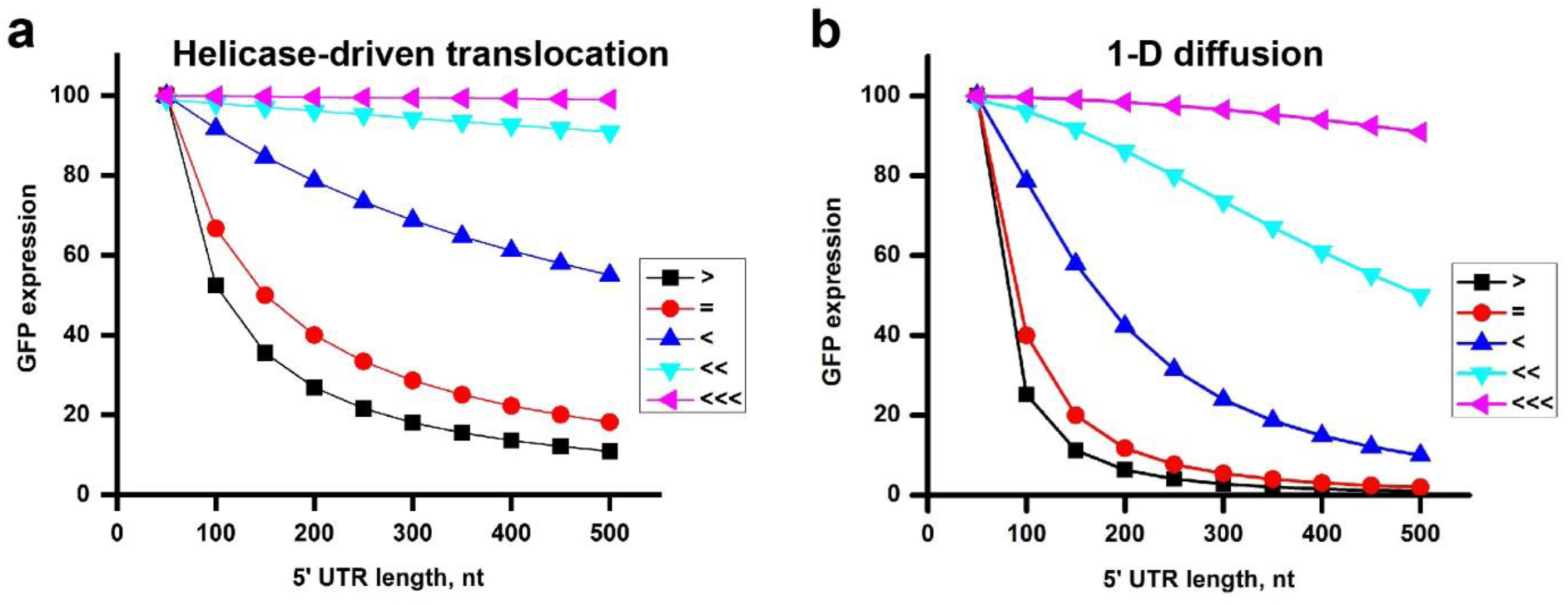
Modeled dependencies of GFP synthesis on the length of the 5′ UTR for the helicase-driven (Eq. 1) and one-dimensional diffusion (Eq. 2) mechanisms of 40S scanning. Amounts of synthesized GFP are inversely proportional to 5′ UTR scanning time and sum of all other steps of initiation, elongation and termination, which do not depend on the 5′ UTR length. GFP synthesis is modeled for scenarios when the sum of all other steps of translation, which are independent of 5′ UTR length, are 10-fold faster (> black), equal to (= red), 10-fold slower (< blue), 100-fold slower (<< cyan) or 1000-fold slower (<<< magenta) than scanning of 50 nt-long 5′ UTR.

Both translocation and 1-D diffusion models predict that if scanning is much faster than the sum of all other steps of protein synthesis, protein synthesis is expected to be nearly independent of 5′ UTR length (Fig. 1). Conversely, in case of 40S scanning being comparable to or even slower than the sum of all other steps of translation, protein synthesis is expected to show strong dependence on the 5′ UTR length (Fig. 1). Under these conditions, it might be possible to distinguish between translocation and diffusion mechanisms of scanning by fitting the dependence of GFP synthesis on the 5′ UTR length to equations 1 and 2. Another prediction that can be inferred from these simplified models is that reduction of scanning rate (e.g. via mutation) is expected to more significantly affect translation of mRNAs with longer 5′ UTRs. On the contrary, slowing down any other step of translation is expected to have a stronger relative effect on translation of mRNAs with shorter 5′ UTRs because scanning-independent steps of translation make larger contributions to the time needed to synthesize one GFP molecule on mRNAs with shorter 5′ UTRs than on mRNAs with longer 5′ UTRs.

### Order-of-magnitude variations in the length of unstructured 5′ UTR have a modest effect on translation in yeast cells

To experimentally test how the length and secondary structure of the 5′ UTR affect translation and which protein factors critical for scanning in live cells, we employed the RNA-ID reporter system (Fig. 2a) (45). The RNA-ID reporter provides a robust and sensitive way to examine the effects of cis-regularity mRNA sequences on translation in yeast cells. The RNA-ID DNA construct contains the ORFs of superfolder GFP and the yeast codon-optimized red fluorescent protein variant of mCherry under control of the bidirectional *GAL1,10* promoter, with the *MET15* or *URA3* gene used for selection in yeast, and flanking sequences allowing for the integration into the *ADE2* locus of the yeast genome via homologous recombination. GFP levels are measured by GFP fluorescence normalized by the fluorescence of RFP to account for differential activation of the promoter in different cells.

**Figure 2.**
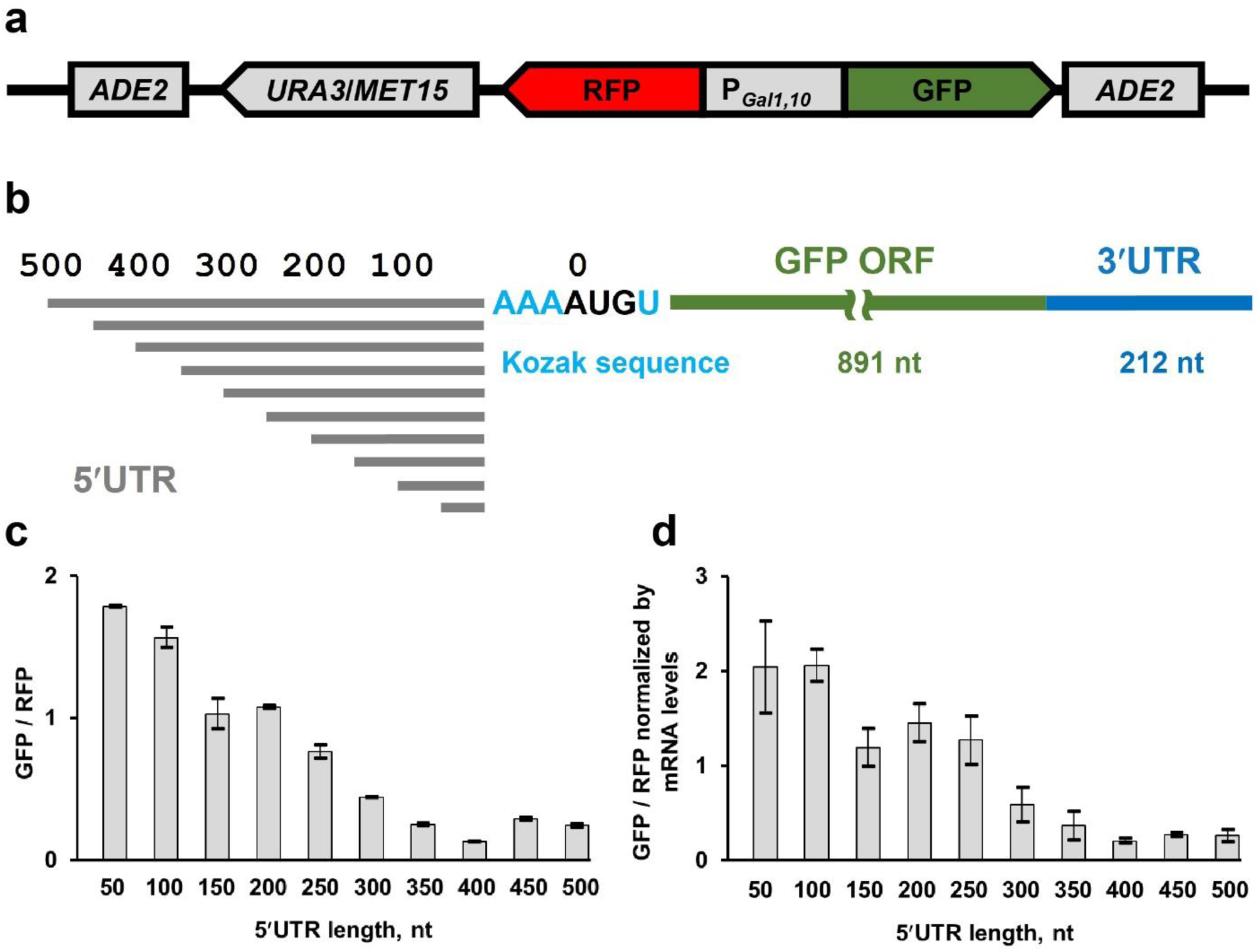
Lengthening unstructured 5′ UTR enriched with UCC and ACCAC sequences progressively reduces translation of GFP reporter in yeast cells. **(a)** Schematic depiction of RNA-ID reporter that includes *ADE2* loci for integration into the yeast genome, *URA3* or *MET15* selection marker gene (details in Materials and Methods), *Gal 1,10* bi-directional promoter, RFP and sfGFP ORFs. **(b)** Design of GFP mRNAs containing unstructured 5′ UTRs (NUS′) of different lengths. The constructs share the same NUS sequence, but differ in how much of the NUS sequence is included from the 5’ end. **(c)** Mean GFP/RFP fluorescence ratios measured in yeast cells, which were transformed with different RNA-ID 5′ UTR NUS′ constructs as indicated. GFP/RFP ratio in GO RNA-ID with 6 nt-long 5′ UTR was set to 1. **(d)** Mean GFP/RFP fluorescence ratios normalized to the ratio of GFP and RFP mRNA levels, which were determined by RT-qPCR. Error bars show standard deviations determined from three biological replicates.

In all RNA-ID constructs, the ORF of superfolder GFP was flanked by the 3′ UTR from yeast RPL41B mRNA (Fig. 1a). RPL41B mRNA is the one of most well-translated mRNAs in yeast cells; the 3′ UTRs of this mRNA were previously precisely identified using rapid amplification of cDNA ends (RACE) (46). To examine the effect of the 5′ UTR length on translation, upstream of the GFP ORF, we introduced intrinsically unstructured sequences ranging from 50 to 500 nucleotides in length with 50 nucleotide increments at the 5′ end of the 5′ UTR while preserving the original Kozak context of the start codon (Supplementary Table1, Fig. 2b). These non-repetitive unstructured sequences (NUS) were designed with the previously developed genetic algorithm (36). The genetic algorithm evolves a sequence through an iterative sequence alteration/selection process that eliminates all alternative basepairing interactions, including those pairs that the 5′ UTR nucleotides can form with the entire mRNA sequence including the ORF and 3′ UTR. A single 500 nucleotide NUS was designed, and individual constructs use differing lengths were created by truncation of the 5′ end. Therefore, these constructs share the sequence upstream of the Kozak sequence and the start codon. Importantly, in contrast to CA and CAA repeats commonly used to eliminate RNA secondary structure, NUSs are more complex and more easily propagated in *E.coli* and yeast cells. Basepairing probability estimated by the RNAstructure software package predicts that NUS nucleotides are nearly completely unpaired while most nucleotides in the ORF and 3′ UTR have high (>75%) basepairing probability (Suppl. Fig. S1).

Resulting RNA-ID constructs (named ^50^NUS′-^500^NUS′ where superscript numbers indicate 5′ UTR length) were transformed into BY4741 haploid strain of *Saccharomyces cerevisiae*. GFP/RFP ratio for each construct was measured by flow cytometry. Extending the 5′ UTR progressively reduced GFP synthesis until it leveled off at the length of 350-400 nucleotides (Fig. 2c). Normalizing GFP/RFP fluorescence by the GFP/RFP ratio of mRNA levels, which were determined by RT-qPCR, indicated that observed reduction in GFP synthesis was due to decrease in translational efficiency rather than changes in mRNA levels (Fig. 2d). Translation of GFP mRNAs with 350-500 nt-long unstructured 5′ UTRs was ∼ 8 times lower on average than translation of mRNA with a 50 nt-long 5′ UTR. These data showed considerable dependence of translation on the 5′ UTR length. Alternatively, observed reduction in GFP levels could be due to sequence-specific effects. Indeed, further analysis of ^50^NUS′-^500^NUS′ sequences revealed that extending the 5 ′UTR from 50 to 500 nt increasingly enriched NUS sequences with C-rich motifs UCC and ACCAC (Suppl. Fig. S2). These C-rich sequences were recently reported to be inhibitory for translation initiation (47).

To test whether observed reduction of protein synthesis in ^50^NUS′-^500^NUS′ constructs was due to the presence of inhibitory C-rich sequences or increased 5′ UTR length, we computationally design a new set of unstructured 5′ UTRs (^50^NUS1-^500^NUS1) ranging in length from 50 to 500 nt, in which UCC and ACCAC sequences motifs were avoided (Supplementary Table 1). Because sequences adjacent to the 5′ cap may influence 40S recruitment, we extended NUS1 UTRs with 50 nucleotide increments at the 3’ end while keeping the 5′ end constant (Fig. 3a). As expected, RFP fluorescence remained nearly constant between strains transformed with different RNA-ID NUS1 constructs while extending the 5′ UTR in GFP mRNA moderately reduced GFP synthesis (Fig. 3 b-d). Transforming ^50^NUS1-^500^NUS1 constructs into BY4741 and BY4742 strains produced similar results (Suppl. Fig. S3). Translation of ^50^NUS1 was only two-fold more efficient than translation of ^500^NUS1 (Fig. 3c-d). Hence, the larger reduction of translational efficiency observed in ^50^NUS′-^500^NUS′ constructs might be due to presence of UCC/ACCAC and, possibly, other inhibitory sequences.

**Figure 3.**
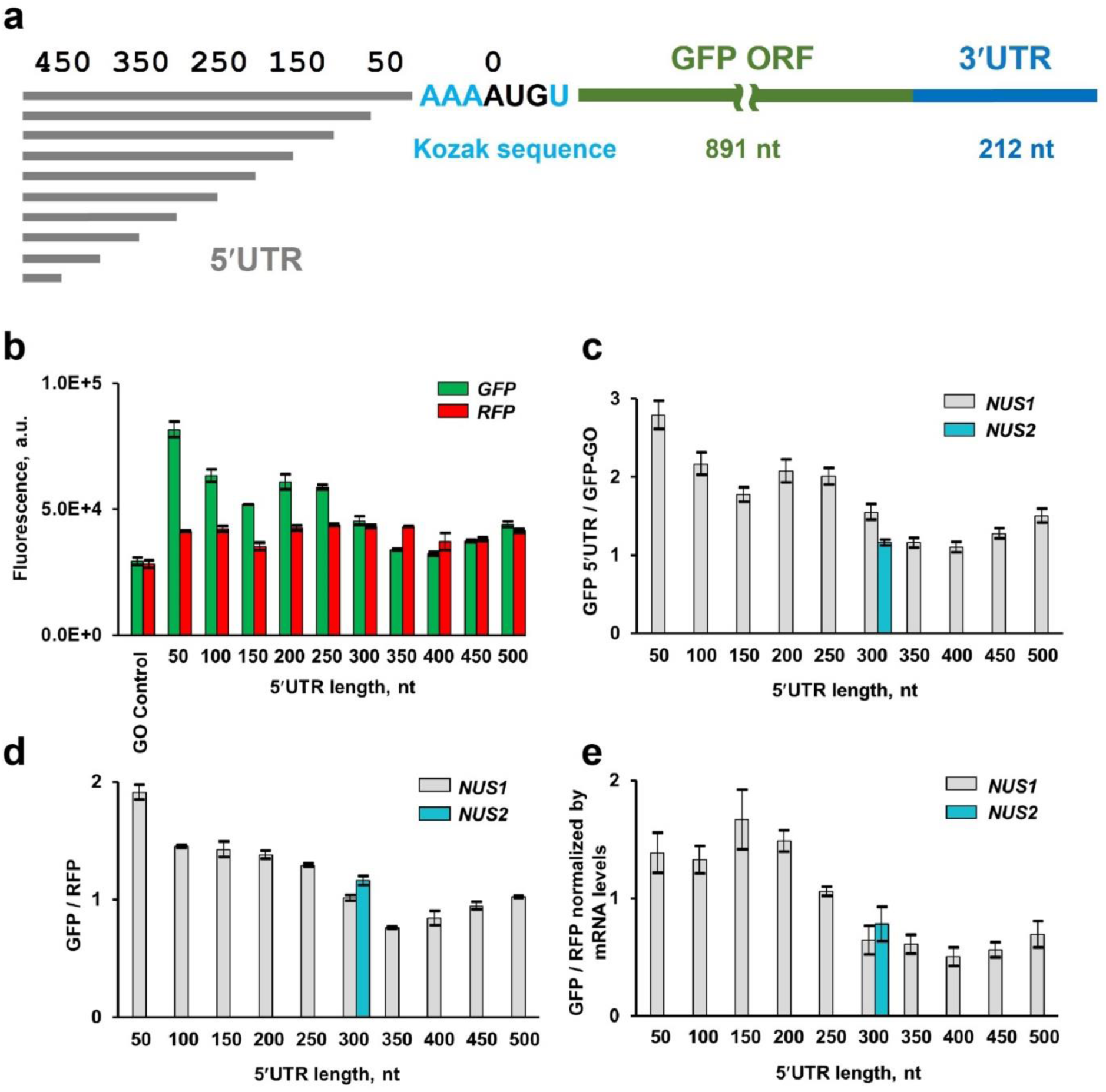
10-fold lengthening of unstructured 5′UTR moderately reduces translation of GFP reporter in yeast cells. **(a)** Design of GFP mRNAs containing unstructured 5′ UTRs (NUS1 series) of different lengths. **(b)** Mean GFP and RFP fluorescence measured in yeast cells, which were transformed with different RNA-ID constructs as indicated. **(c)** Mean GFP fluorescence in strains expressing RNA-ID mRNA with unstructured 5′ UTRs of different lengths normalized by GFP fluorescence produced in GO RNA-ID control reporter with 6 nt-long 5′ UTR. **(d)** Mean GFP/RFP fluorescence ratios. GFP/RFP ratio in GO RNA-ID control was set to 1. **(e)** Mean GFP/RFP fluorescence ratios normalized to the ratio of GFP and RFP mRNA levels, which were determined by RT-qPCR. NUS1 and NUS2 are independently designed, divergent non-repetitive unstructured sequences. Error bars show standard deviations determined from three biological replicates.

To further explore how sequence variations in the unstructured 5′UTR affect translation, we independently designed another 300 nt-long 5′UTR (^300^NUS2), in which UCC and ACCAC sequences were also avoided. ^300^NUS1 and ^300^NUS2 shared only 57.3% identity in a global pairwise alignment calculated by the EMBOSS Stretcher alignment tool and 58.8% identity in a local alignment calculated by the EMBOSS Water alignment tool (48). Similar sequence identity (mean of 56.9% and standard deviation 2.1%) was found in local alignment of 100 computationally shuffled (i.e. randomized) ^300^NUS1 and ^300^NUS2 sequences. GFP/RFP ratio in ^300^NUS1 and ^300^NUS2 were similar (Fig. 3c-d) indicating that as long as inhibitory motifs were eliminated from the design, other sequence variations had minimal effect on translation. We thus focused the rest of our experiments on ^50^NUS1-^500^NUS1 constructs.

Normalizing GFP/RFP ratio by mRNA levels, which were determined by RT-qPCR, showed that observed changes in GFP synthesis in ^50^NUS1-^500^NUS1 strains were due to modest reduction of translational efficiency rather than changes in mRNA levels (Fig. 3e). It is noteworthy that GFP/RFP ratios (Fig. d-e) and GFP levels alone (Fig. 2c) showed similar dependence of 5′ UTR length. Hence, transcriptional activity of *Gal1,10* promoter is effectively constant in different cells, and GFP fluorescence alone (without normalizing to RFP levels) can be reliably used to analyze changes in translational efficiency of GFP reporter constructs.

To test whether in yeast cells, ^50^NUS1-^500^NUS1 mRNAs are transcribed from the previously identified transcription start site of the *GAL1,10* promoter, we performed 5′ RACE analysis for ^50^NUS1, ^300^NUS1 and ^500^NUS1 mRNAs. 5′ RACE PCR products were of expected size and sequence (Suppl Fig. S4a) consistent with the predicted transcription start. SDS-PAGE analysis combined with GFP fluorescence imaging revealed a single fluorescent GFP protein band in all ^50^NUS1-^500^NUS1 strains indicating that only one ORF, which produced fluorescent GFP, was predominately translated (Suppl. Fig. S4b). Hence, no alternative near-cognate, in-frame start sites were used for translation initiation in ^50^NUS1-^500^NUS1 GFP mRNAs.

### RNA secondary structures in the 5′UTR strongly inhibit translation

Our experiments with ^50^NUS1-^500^NUS1 reporter constructs showed that extending the 5′ UTR 10-fold over the median length of yeast 5′ UTRs only modestly reduces protein synthesis. Therefore, 40S scanning along unstructured 5′ UTR seems quite effective and does not hamper translation initiation. We next explored the effects of 5′ UTR secondary structure on translational efficiency and 40S scanning. To that end, we replaced nucleotides 156-199 of ^300^NUS2 5′UTR with a GC-rich stem-loop containing 20 basepairs and a UUAA tetraloop (Fig. 4a). When compared to the ^300^NUS2 5′ UTR, the ^300^NUS2 5′ UTR with the stem-loop (^300^NUS2-SL) produced ∼150-fold less GFP (Fig. 4b). When adjusted by mRNA levels measured by RT-qPCR, the stem-loop reduced GFP translation by 85-fold (Fig. 4c). Likewise, we replaced 44 nucleotides 150 nt downstream of the 5′ end of ^300^NUS′ and ^500^NUS′ 5′ UTRs with a stem-loop containing 20 base pairs and a tetraloop. The insertion of this stem-loop into ^300^NUS′ and ^500^NUS′ 5′ UTRs inhibited GFP synthesis by 30- and 10-fold, respectively (Suppl. Fig. S5). Hence, consistent with many published reports (11–15), stable RNA secondary structure strongly inhibit translation, presumably by hindering 40S scanning. These results also indicate that ^50^NUS-^500^NUS (as well as ^50^NUS′-^500^NUS′) mRNA constructs are translated in cap- and scanning-dependent manner rather than through a non-canonical, cap-independent mechanisms (e.g. IRES-dependent initiation).

**Figure 4.**
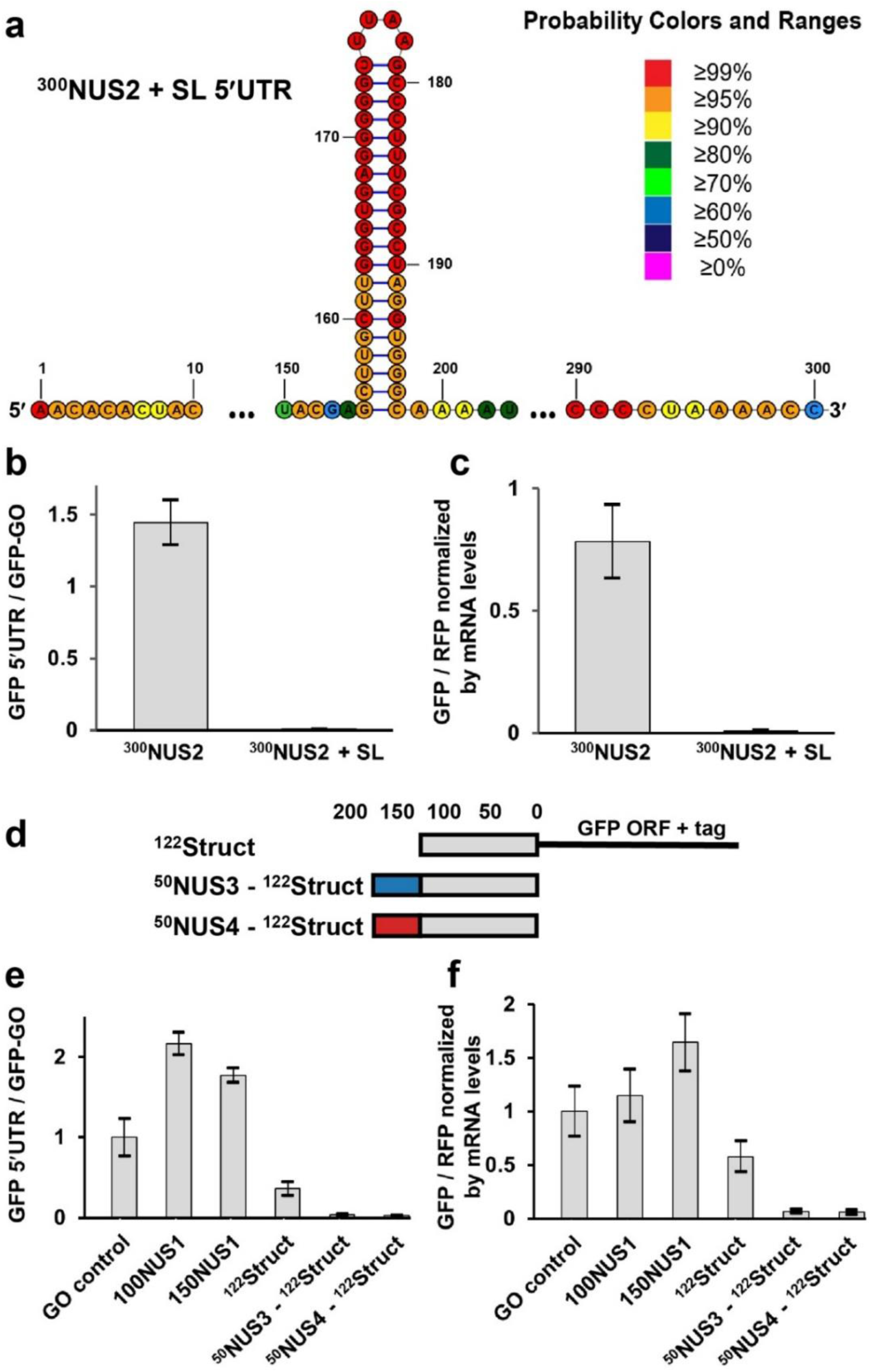
RNA secondary structure in the 5′UTR strongly inhibits translation. **(a)** 54 nucleotides in the middle of ^300^NUS2 5′UTR were replaced with 20-bs long RNA stem-loop to create ^300^NUS2+SL mRNA. Basepair probabilities determined by *RNAstructure* software is shown by color code as indicated. **(b-c)** Introducing the stem-loop (SL) into ^300^NUS2 5′ UTR reduced GFP synthesis. **(d-f)** 122-nt long sequence (^122^Struct) in the 5′ UTR (grey box in schematics **d**), which is predicted to fold into multiple stem-loops, inhibits GFP synthesis. **(b, e)** Mean GFP fluorescence was normalized to that in GO RNA-ID reporter with 6 nt-long 5′ UTR. **(c, f)** Mean GFP/RFP fluorescence ratios normalized to the ratio of GFP and RFP mRNA levels, which were determined by RT-qPCR. GFP/RFP ratio in GO RNA-ID control was set to 1. Error bars (**b-c, e-f**) show standard deviations determined from three biological replicates.

Next, instead of introducing a very stable stem-loop into an unstructured 5′ UTR, we replaced the entire 5′ UTR with a 122-nt long sequence that is derived from the 5′ UTR of human GAPDH mRNA to make the ^122^Struct construct. The RNAstructure software package predicts that most nucleotides of this sequence have between 50 and 75% probability of basepairing (Suppl. Fig. 6-7). Hence, rather than forming one very stable secondary structure, the ^122^Struct sequence likely folds into a dynamic ensemble of alternative structures. When compared to ^100^NUS1 and ^150^NUS1 strains, ^122^Struct strain produced 5- and 6-fold less GFP, respectively (Fig. 4e). The GFP/RFP ratio normalized to mRNA levels in ^122^Struct strain was 2 and 3-fold lower than that in ^100^NUS1 and ^150^NUS1 strains, respectively (Fig. 4f), indicating that a structured 5′ UTR substantially reduces translation relative to unstructured 5′ UTR of similar length.

Because the ^122^Struct 5′ UTR is predicted to contain basepairs near the 5′ end (Suppl. Fig. S6-S7), secondary structures of the ^122^Struct 5′ UTR may inhibit both initial 40S recruitment and following 40S scanning (11). We next aimed to eliminate the effects of secondary structure on 40S recruitment. As the 43S initiation complex covers ∼45 nucleotides of mRNA (49,50), we added 50 nt-long unstructured sequences NUS3 or NUS4 upstream of ^122^Struct sequence that would allow for unhindered 40S binding. Surprisingly, adding either NUS3 or NUS4 unstructured sequence reduced GFP levels 10- or 14-fold, respectively, relative to GFP levels in ^122^Struct 5′ UTR strain (Fig. 4e). GFP/RFP ratios normalized to mRNA levels in NUS3-^122^Struct and NUS4-^122^Struct strains were 9-fold lower than that in ^122^Struct 5′ UTR strain (Fig. 4f). One possible explanation of these results is that 40S binding to the original ^122^Struct mRNA partially unwound secondary structures of the 5’ UTR and thus aided 40S scanning. 40S recruitment to the unstructured sequence (NUS3 or NUS4) upstream of the ^122^Struct preserved all secondary structure elements within^122^Struc sequence and thus reduced effectiveness of 40S scanning on NUS3-^122^Struct and NUS4-^122^Struct mRNAs. Taken together, experiments with ^300^NUS2-SL and ^122^Struct mRNAs demonstrate that secondary structure elements within 5′ UTR strongly inhibit protein synthesis by impeding 40S scanning.

### Translational helicase eIF4A is not involved in 40S movement along the 5′ UTR

We next used mutants to test whether mRNAs with long unstructured 5′ UTRs are particularly sensitive to genetic perturbations of translation factors, which were previously implicated in 40S scanning. DEAD-box helicase eIF4A is a subunit of 5′ cap recognition complex eIF4F. Helicase activity of eIF4A is thought to be directly involved in ATP-dependent 40S movement along 5′ UTR (21,51,52). In yeast, eIF4A is encoded by two paralogues *TIF1* and *TIF2*. To test involvement of eIF4A in 40S scanning, RNA-ID constructs were transformed into either into *tif1Δ* or *tif2Δ* deletion strains created in BY4741 genetic background. RT-qPCR measurements showed ∼two-fold decrease in eIF4A mRNA levels in both *tif1*Δ and *tif2*Δ strains when compared to the parent (“wild-type”) BY4741 strain (Suppl Fig. S8). Nevertheless, GFP synthesis on ^50-500^NUS1 and ^122^Struct mRNAs were essentially unaffected by either *tif1* or *tif2* deletion (Fig. 5a). Likewise, GFP synthesis on ^50-500^NUS′ mRNAs were unchanged in *tif1Δ* or *tif2Δ* strains (Suppl. Fig. S9a). This could be due to high expression levels of eIF4A, which is the most abundant factor of translation initiation in yeast cells (53). Consistent with the observation that protein synthesis was unaffected by deletion of either *tif1* or *tif2*, growth of *tif1Δ* or *tif2Δ* strains on YPD plates was similar to the parental, wild-type (BY4741) strain (Suppl. Fig. S10).

**Figure 5.**
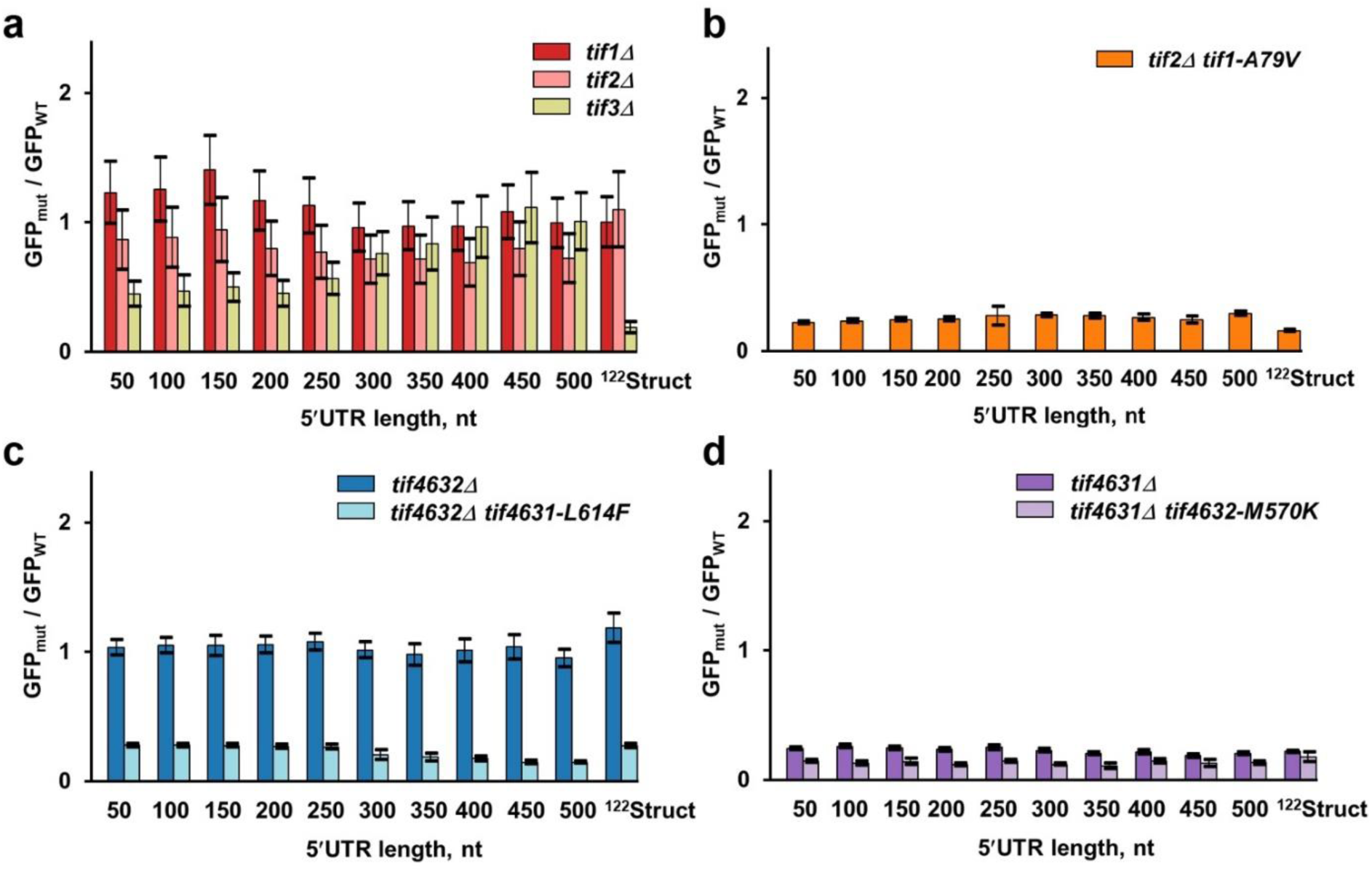
Mutational perturbations of eIF4F subunits similarly reduce translation of GFP mRNAs with short and long unstructured 5′ UTRs. GFP fluorescence measured in mutant strains normalized to respective GFP fluorescence measured in wild-type strain. Error bars indicate standard deviations determined from three biological replicates. (**a**) eIF4A-encoding *tif1, tif2* or eIF4B-encoding *tif3* were deleted in BY4741 background as indicated. **(b)** Either wild-type (*TIF1*) or temperature sensitive (*tif1-A79V*) alleles of eIF4A-encoding *tif1* were expressed in the strain lacking both chromosomal copies of *tif1* and *tif2*. **(c)** An amino acid substitution L614F, which reduces eIF4G binding to eIF4A, was introduced into *tif4631* in the background of *tif4632* deletion to create temperature-sensitive *tif4632*Δ*tif4631-L614F* strain. **(d)** An amino acid substitution M570K, which reduces eIF4G binding to eIF4A, was introduced into *tif4632* in the background of *tif4631* deletion to create temperature-sensitive *tif4631*Δ*tif4632-M570K* strain. BY4741 strain is considered as wild-type in panels **a, c** and **d.** In experiments with temperature-sensitive strains *tif2*Δ*tif1-A79V* and *tif4632*Δ*tif4631-L614F, Gal 1,10* promoter was induced concurrently with raising incubation temperature from 30 to 37°C degree. Respective wild-type control strains received identical treatment.

Because eIF4A is essential for yeast viability we could not delete both *TIF1* and *TIF2* genes simultaneously. Instead, we integrated RNA-ID constructs into the strain lacking both *TIF1 and TIF2* genes and expressing the temperature sensitive allele *tif1-A79V* (31). A79V substitution is thought to weaken eIF4A binding to RNA (31,54,55) and severely hamper yeast growth at 37°C (Suppl. Fig. S11). To examine effects of temperature sensitive A79V mutation in eIF4A, GFP expression was induced by addition of galactose concurrently with switching from 30 to 37°C. In the absence of galactose, negligible levels of GFP were accumulated, indicating tight transcriptional control of the *Gal 1,10* promoter (Suppl. Fig. S12). When compared at 37°C to the strain, in which wild-type *TIF1* allele was expressed, expressing *tif1-V79A* decreased GFP levels 4-fold in all ^50-500^NUS1 constructs irrespective of 5′ UTR length (Fig. 5b). Similar uniform reduction of GFP synthesis was observed for all ^50-500^NUS′ mRNAs (Suppl. Fig. S9b). Translation on the ^122^Struct mRNA containing a 122 nt long structured 5′ UTR was reduced 6-fold, i.e. fairly similar to translation of mRNAs with unstructured 5′ UTR (Fig. 5b). This result is consistent with published data indicating that eIF4A stimulates translation of mRNAs regardless of structural complexity of their 5′ UTR (56).

Helicase activity of eIF4A is stimulated by another initiation factor, eIF4B, which is encoded by a single-copy gene *TIF3*. We next deleted non-essential gene *TIF3* in a BY4741 background. In the *tif3*Δ strain, translation of GFP mRNAs with 50-200 nt long unstructured 5′ UTRs was reduced two-fold, while translation of GFP mRNAs with 300-500 nt long unstructured 5′ UTRs was unaffected (Fig. 5a). Similar results were obtained with ^50-500^NUS′ reporters (Suppl. Fig. 9a). Remarkably, the ^122^Struct mRNA, containing a 122 nt long structured 5′ UTR, was far more sensitive to eIF4B deletion as GFP levels decreased 6-fold in the *tif3*Δ strain. This observation agrees with ribosome profiling data showing that eIF4B stimulates translation of mRNAs with structured 5′ UTRs (57).

eIF4A helicase and ATPase activities are stimulated by interaction with another subunit of eIF4F complex, eIF4G (58–60). In yeast, eIF4G is encoded by two paralogues, *TIF4631* and *TIF4632.* When we deleted the *TIF4632* gene, translation of all RNA-ID constructs was unaffected (Fig. 5c). We next introduced an amino acid substitution L614F into *tif4631* in the presence of *tif4632Δ*. L614F mutation weakens interaction of eIF4G with eIF4A and produces a temperature sensitive phenotype, which can be suppressed by overexpression of eIF4A or eIF4B (61) (Suppl. Fig. S11). When measured at 37°C, the L614F mutation in *tif4631* reduced GFP synthesis of ^50^NUS1, ^500^NUS1 and ^122^Struct mRNAs 4-, 7- and 4-fold, respectively (Fig. 5c). Relative to *tif4632Δ* strain, GFP levels in *tif4632Δ tif4631-L614F* strain transformed with either ^50^NUS′ or ^500^NUS′ RNA-ID reporters were equally reduced by 3-fold (Suppl. Fig. 9c).

We also made reciprocal mutational perturbations in *tif4632* and *tif4631.* In that, we deleted *tif4631* and introduced an amino acid substitution M570K, which weakens interaction of eIF4A with eIF4G (61), into *tif4632.* Deletion of *tif4631* lowered GFP levels 5-fold irrespective of 5′ UTR length in both ^50-500^NUS1 and ^50-500^NUS′ reporters (Fig. 5d, Suppl. Fig. S9d). Mutation M570K in *tif4632* further reduced GFP levels 1.5-2-fold in all 5′ UTR NUS1 and NUS′ reporters (Fig. 5d, Suppl. Fig. S9d). Relative to the *tif4631Δ* strain, translation of the ^122^Struct mRNA decreased by 1.2-fold in the *tif4631Δ tif4632-M570K* strain (Fig. 5d).

Taken together, our results indicate that amino acid substitutions disrupting eIF4A-eIF4G interaction similarly affected GFP reporters with short and long of 5′ UTRs. Hence, the eIF4F complex as well as eIF4A alone are unlikely to be responsible for 40S movement along the 5′ UTR.

### Loss-of-function mutations in eIF3g and eIF3i moderately affect translation of mRNAs with short and long unstructured 5′ UTRs

We next probed whether initiation factors eIF3g and eIF3i, which are encoded by yeast genes *TIF35* and *TIF34*, respectively, affect 40S scanning. These two subunits of multimeric initiation factor 3 bind near the mRNA entry site of the 40S subunit, i.e. the “front” side of the small subunit in respect to the direction of scanning. eIF3g and eIF3i were shown to stimulate ATPase activity of eIF4A (56). In a recent cryo-EM reconstruction of the 48S initiation complex, two eIF4A molecules were visualized. One was bound at the mRNA entry site; the other was bound to the eIF4G•eIF4E•5′cap near the “rear” side of the small subunit (52). eIF3i was seen making extensive interactions with eIF4A bound to the mRNA entry site, providing further evidence for possible eIF3i involvement in helicase and translocase activities of the 43S initiation complex.

In yeast, triple alanine substitution K194A/L235A/F237A of conserved residues in the RRM of eIF3g (*tif35-KLF*) or a single-point mutation Q258R in the WD40 repeat 6 of eIF3i (*tif34-Q258R*) were shown to produce growth defects (62). These mutations also affected efficiency of re-initiation on *GCN4* mRNA, which contains four short uORFs upstream of the main ORF (62). Based on these observations, it was hypothesized that eIF3g and eIF3i may be involved in 40S scanning (62). We re-created these eIF3g and eIF3i mutations in the BY4741 genetic background (Suppl. Fig. S11). When compared to BY4741, in the temperature-sensitive *tif35-KLF* strain, GFP expression decreased 2-2.5-fold in all ^50-500^NUS1 RNA-ID constructs irrespective of 5′ UTR length (Fig. 6a), showing no evidence of eIF3g involvement in 40S scanning. Translation on the ^122^Struct mRNA was reduced 1.5-fold, similar to translation of mRNAs with unstructured 5′ UTRs (Fig. 6a).

**Figure 6.**
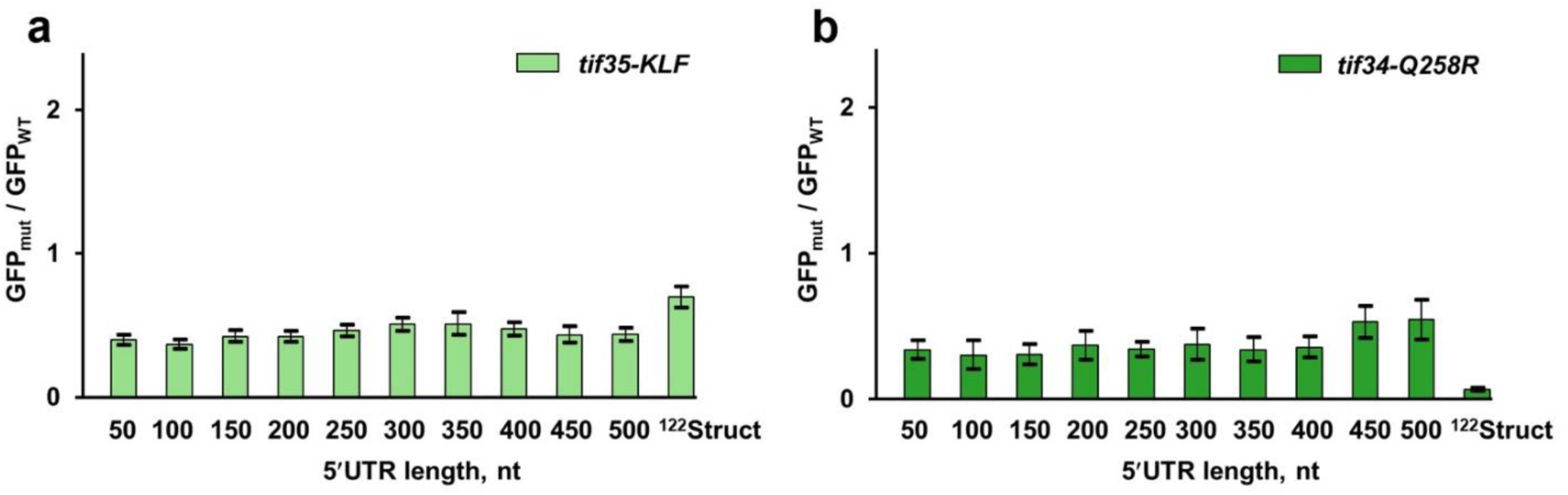
Mutations in eIF3g and eIF3i similarly reduce translation of GFP mRNAs with short and long unstructured 5′ UTRs. Amino acid substitutions were introduced in eIF3g-encoding *tif35* (**a**) or eIF3i-encoding *tif34* (**b**) in the BY4741 genetic background. GFP fluorescence measured in mutant strains was normalized to respective GFP fluorescence measured in wild-type (BY4741) strain. Error bars indicate standard deviations determined from three biological replicates. In experiments with temperature-sensitive *tif35-KLF* strain*, Gal 1,10* promoter was induced concurrently with raising incubation temperature from 30 to 37°C degree for both *tif35-KLF* and wild-type cells.

Amino acid substitution Q258R in eIF3i affected translation of GFP mRNAs with shorter unstructured 5′ UTRs somewhat more significantly than translation of mRNA with longer 5′ UTRs: GFP expression on ^50^NUS1 and ^500^NUS1 mRNAs decreased 4 and 2 times, respectively (Fig. 6b). In contrast to moderate reduction of GFP synthesis on mRNAs with unstructured 5′ UTRs, Q258R substitution in eIF3i diminished translation of ^122^Struct mRNA 15-fold (Fig. 6b). Hence, eIF3i may be involved in unwinding of secondary structure in the 5′ UTRs rather than 40S movement along mRNA itself.

### Translational helicases Ded1 and Slh1 are not involved in 40S movement along the 5′ UTR

We next examined involvement of two translational helicases Ded1 and Slh1 in 40S movement along the 5′ UTR. Ded1 (yeast ortholog of mammalian DDX3X) is a DEAD box helicase that is implicated in unwinding stable stem-loops in 5′ UTRs during translation initiation (31,63). We transformed RNA-ID constructs into a yeast strain containing the cold sensitive allele *ded1-120* (Suppl. Fig. S11), which encodes Ded1 bearing amino acid substitutions G108D and G494D (31). These mutations are thought to impair ATP binding or hydrolysis (54). GFP expression in the cold-sensitive *ded1-120* strain (Suppl. Fig. S11) was induced by addition galactose simultaneously with the temperature shift from 30 to 18°C. When compared to GFP expression in BY4741 strain at 18°C, translation of all reporter mRNAs with unstructured 5′ UTRs in *ded1-120* strain was reduced (Fig. 7a). Contrary to the idea of Ded1 involvement in 40S translocation during scanning (32), mRNAs with shorter unstructured 5′ UTRs were more significantly affected as GFP expression on ^50^NUS1 and ^500^NUS1 mRNAs decreased 7 and 3-fold, respectively (Fig. 7a). Similar results were obtained with ^50-500^NUS′ reporters (Suppl. Fig. S9e). Higher relative sensitivity of mRNAs with shorter 5′ UTRs to Ded1 mutation are consistent with the idea that Ded1 affects scanning-independent steps of translation such as 40S recruitment (23). Slowing down scanning-independent steps of initiation make larger contributions to the time needed to synthesize one GFP molecule on mRNAs with shorter 5′ UTRs (Equations 1-2).

**Figure 7.**
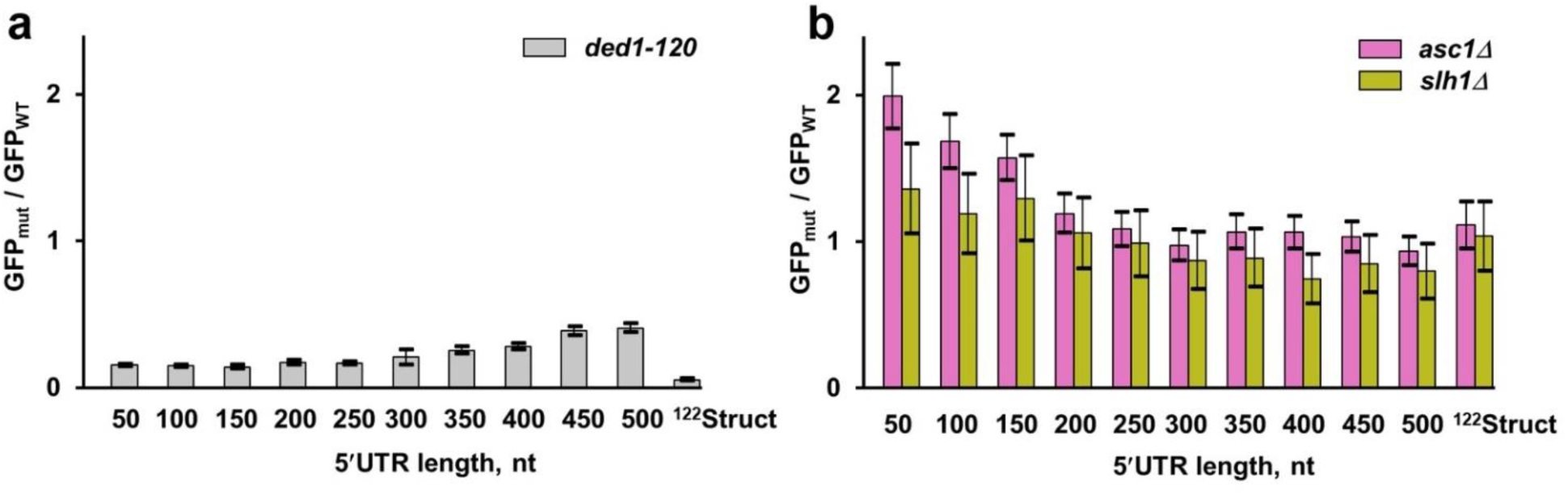
Translational helicases Ded1 and Slh1 are not essential for 40S scanning. GFP fluorescence measured in mutant strains normalized to respective GFP fluorescence in wild-type (BY4741) strain. Error bars indicate standard deviations determined from three biological replicates. (**a**) RNA-ID constructs were transformed into a *ded1-120* strain containing cold-sensitive allele *ded1-120*, which encodes mutant form of Ded1 helicase baring amino acid substitutions G108D and G494D. (**b**) RNA-ID constructs were transformed into *slh1*Δ or *asc1*Δ strains, which helicase Slh1 and ribosomal protein Asc1, respectively. In panels **a** GFP synthesis was induced concurrently with temperature shift from 30 to 17°C.

Consistent with the importance of Ded1-mediated unwinding of mRNA secondary structure (23,31,63), translation of the ^122^Struct mRNA containing a structured 5′ UTR diminished nearly 20-fold in *ded1-120* cells (Fig. 7a). Taking together, our data support the model suggesting that Ded1 is critical for melting secondary structures but not the actual movement of the 40S along the 5′ UTR.

Recently, another translational helicase, ski2-like ASCC3, was linked to 40S scanning, as ASCC3 knockdown lowered translational efficiency and 40S occupancy along the 5′ UTR in mammalian cells (64). ASCC3 and its yeast orthologue Slh1 are known to disassemble stalled 80S ribosomes in ribosome-associated quality control (RQC) pathway (65). Noteworthy, ASCC3 was shown to be 3′-to-5′ translocase, i.e. it moves in the direction opposite to scanning 40S subunit (66,67). Nevertheless, we tested whether yeast Slh1 may be responsible for the 40S movement along the 5′ UTR by deleting the single non-essential gene encoding this helicase. Because Slh1 interacts with non-essential protein Asc1 on the small ribosomal subunit (67), we also expressed GFP reporters in *asc1Δ* strain. Relative to BY4741 wild-type cells, GFP expression of the ^50^NUS1 construct increased 1.4 and 2-fold in *slh1*Δ and *asc1*Δ strains, respectively (Fig. 7b). Neither genetic perturbation substantially affected GFP expression with long 5′ UTRs. Deletions of slh1 and asc1 similarly affected ^50-500^NUS′ reporters (Suppl. Fig. 9f). Hence, our data provide no supporting evidence for Slh1 involvement in 40S scanning.

## Discussion

Our experiments indicate that extending the unstructured 5′ UTR 10-fold over the median length in yeast transcripts reduces GFP synthesis by only ∼2-fold (Fig. 3). Hence, 40S scanning is not rate limiting in yeast cells under normal growth conditions. Our results are consistent with two other studies examining 5′ UTR length dependence of translational efficiency in yeast cells (31,32). In particular, luciferase reporters containing unstructured CA repeats in the 5′ UTR were used to show that extending CA repeats up to 187 nucleotides (4-fold over median the median length of 5′ UTR length in yeast) did not affect luciferase synthesis (31). Several other studies, which showed that lengthening 5′ UTR substantially reduces translation, employed structured sequences (29,30,32,34) and followed protein synthesis in cell extracts rather than in intact cells (29,32).

In contrast to unstructured sequences, elements of secondary structure in the 5′ UTR were shown to inhibit translation in our (Fig. 3) and other studies (11–15). Based on these data, we hypothesize that 5′ UTRs are relatively short in yeast and other taxonomic classes of eukaryotes because lengthening of 5′ UTR sequences increases the number of basepairs and thus hinders 40S scanning. Hence, the extent of secondary structure likely causes negative correlation between 5′ UTR length and gene expression levels (28).

Using RNA-ID reporters and mutagenesis, we also attempted to identify translation factors essential for scanning. The rationale for these experiments was based on several studies indicating that 40S scanning is driven by a processive translocase (20,29,32). Scanning time, which was either measured in single-molecule experiments (20) or inferred from reporter translation in cell extracts (29,32), was observed to scale linearly with 5′ UTR length, which is consistent with helicase-driven translocation of the 40S subunit during scanning. It is noteworthy that assessments of the 40S scanning rate drastically differed between these reports from 6-10 nt/s determined from reporter translation in cell extracts (29,32) to 100 nt/s measured in single-molecule experiments (20). In addition, all three studies employed natural 5′ UTR sequences that have significant propensities to form extensive secondary structures (Suppl. Fig. S13). Hence, in these experiments, 43S movement along 5′ UTRs was likely hindered by and conflated with unwinding of 5′ UTR secondary structures. Furthermore, in single-molecule measurements, scanning rate estimates were based on just four 5′ UTR sequences, which were between 60 and 241 nts in length (20). The relatively small number of points and fairly narrow range of lengths might complicate distinguishing between linear versus parabolic dependencies of scanning time on 5′ UTR length indicative of translocation-driven versus diffusion-based mechanisms of 40S scanning.

In contrast to previous studies of 40S scanning, we employed unstructured 5′ UTR sequences. Loss-of-function mutations or deletion of eIF4B, eIF4G, eIF3g, eIF3i, Asc1 as well as translational helicases eIF4A, Ded1 and Slh1, similarly affected translation of GFP reporters with short and long unstructured 5′ UTRs, indicating lack of involvement of these factors in 40S scanning (Fig. 5-7). These data are consistent with several lines of evidence showing that helicases eIF4A and Ded1 do not drive 40S scanning. (i) Previous studies showed that eIF4A affects mRNA translation irrespective of 5′ UTR length and secondary structure (23,31,32,56). (ii) Experiments performed in a mammalian system reconstituted from purified components demonstrated that the 43S initiation complex can bind to the 5′ end of mRNA and migrate toward the start codon in the absence of eIF4A (68,69). (iii) Single-molecule measurements in a yeast system reconstituted from purified components showed that 40S scanning rate does not depend on concentrations of eIF4A and ATP (20). (iv) A number of biochemical experiments indicate that DEAD box helicases eIF4A and Ded1 do not translocate along RNA and unwind RNA secondary structure by local strand separation with limited processivity (70,71).

Although our data neither rule out participation of eIF4A and Ded1 nor exclude involvement of an unidentified helicase/translocase in 40S movement along 5′ UTR, a parsimonious explanation of our results is that 40S scanning is a helicase-independent, diffusion-based process.

Many experimental findings demonstrate that numerous DNA and RNA binding proteins and protein complexes search for their binding sites via one-dimensional (1-D) diffusion as they slide or hop along DNA/RNA (72–74). When compared to 3-D diffusion, 1-D diffusion greatly accelerates the rate of locating the correct binding site (75–78). 40S scanning might also be driven by 1-D diffusion. Although 1-D diffusion is bi-directional by nature, 40S diffusion along the 5′ UTR may be biased toward the start codon simply by virtue of 40S being recruited to the cap at the 5′ end of mRNA. In that, the 5′ end of mRNA provides a boundary for 1-D diffusion at the beginning of scanning.

Several lines of evidence support the idea that 40S moves along the 5′ UTR by 1-D diffusion. Strong inhibition of translation initiation by 5′ UTR secondary structure observed in our experiments and many published reports (11–15) suggests that 43S initiation complex does not contain a highly-processive, translocating helicase. 3′-to-5′ back-tracking of the 43S initiation complex during scanning was directly observed in *in vitro* single-molecule microscopy experiments (20). AUG triplets placed downstream of the start codon were shown to inhibit translation initiation indicating bi-directional movement of the 43S initiation complex (24). Transcriptome-wide ribosome profiling experiments provide additional evidence for bi-directional scanning and back-sliding of the 40S subunit during translation initiation on mRNA with 5′ UTRs, which are shorter than the 45-nt long footprint of the 43S initiation complex (79). Although further investigation is required, it is likely that 1-D diffusion is, at least in part, responsible for 40S migration from the 5′ cap to the start codon in eukaryotes.

## ACKNOWLEDGEMENTS

We thank Alan Hinnebusch for providing strains H5120-NSY20 (*tif1*Δ *tif2*Δ *tif1-A79V*) and H5121-NSY21 (*tif1*Δ *tif2*Δ*TIF1*) as well as plasmids pEP245 (*tif4631-L614F*) and pEP258 (*tif4632-M570K*). We thank Alan Hinnebusch and Susan Welte for plasmid p5466 (*ded1-120::LEU2*). We thank Eric Phizicky for strains from the single-gene deletion collection and advice on yeast genetics and Christine Lai for her contribution to pilot RNA-ID experiments. We thank Paul Whitford and Thomas Ossevoort for helpful discussions. This work was supported by grants of the National Institutes of Health R35GM141812 (to D.N.E.) and R35GM145283 (to D.H.M.). Phosphoimaging was done using Typhoon RGB instrument, purchase of which was supported by NIH Equipment Grant (S10-OD021489-01A1).

## AUTHORS CONTRIBUTIONS

H.W. and D.N.E. conceived the project. H.W., M.Z., E.G., D.H.M. and D.N.E. designed experiments. H.W. performed experiments. M.Z. designed 5′ UTR sequences and performed computational studies and was supervised by D.H.M. D.N.E. wrote the original draft of the manuscript. H.W., M.Z., E.G, D.H.M. and D.N.E. revised the manuscript.

