## Supplementary Materials for "40S ribosomal subunits scan mRNA for the start codon by one-dimensional diffusion"

### Supplementary Figures

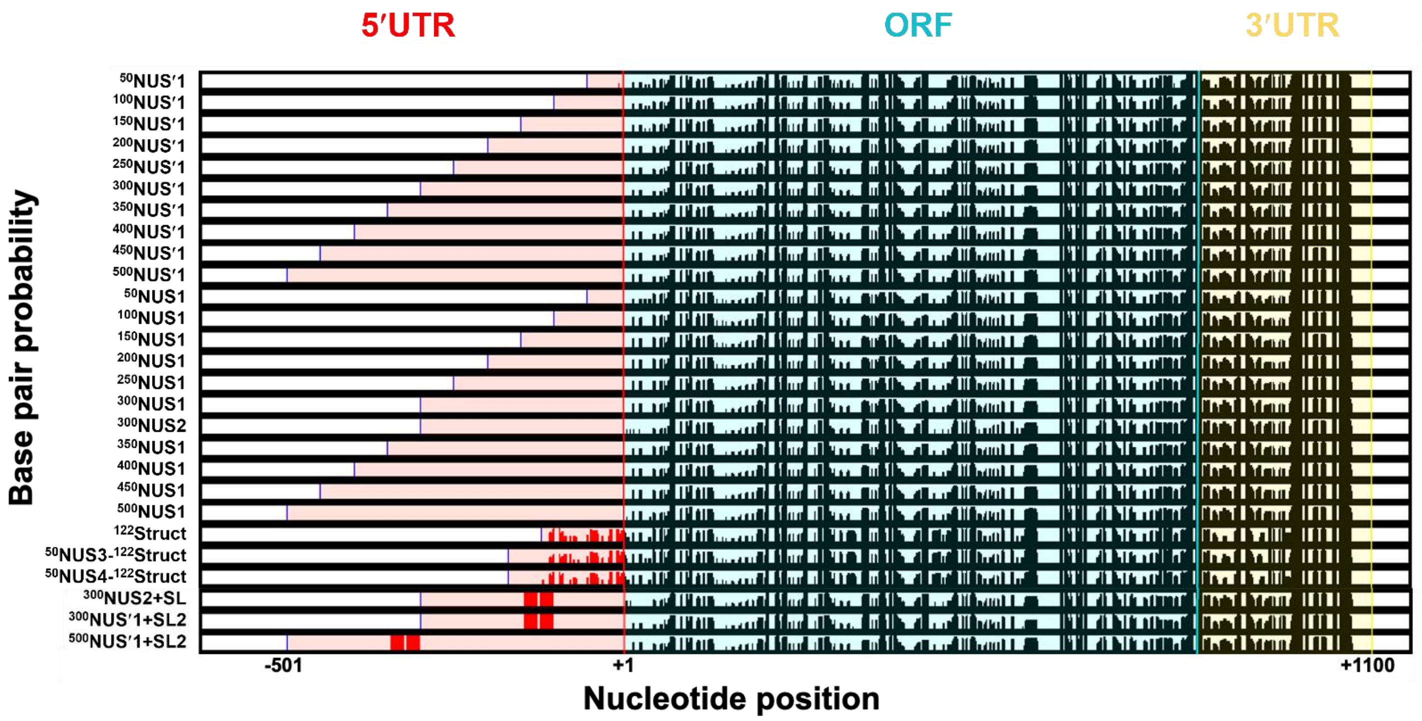


**Figure S1. The nucleotide base pairing probability of GFP mRNA variants.** The base pairing probabilities were estimated by a partition function calculation. The Y-axis is the probability of base pairing. The X-axis represents the nucleotide position in the mRNA. The 5′ UTR, ORF and 3′ UTR are highlighted in red, cyan and yellow, respectively.

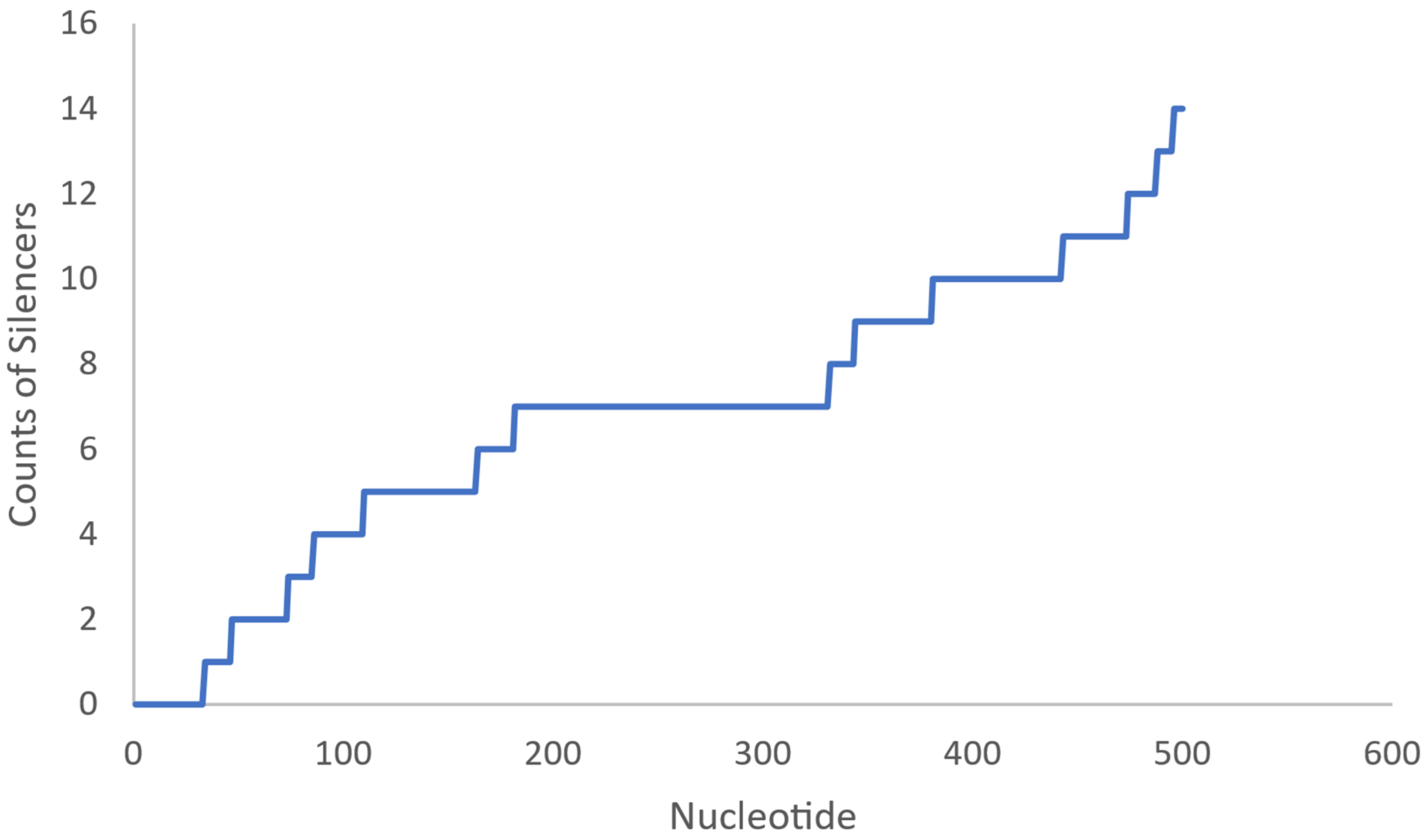


**Figure S2. Cumulative number of UCC and ACCAC translation silencing sequences (y axis) in $^{500}$NUS' 5' UTR (nucleotide position in the $^{500}$NUS' 5' UTR, x axis).**

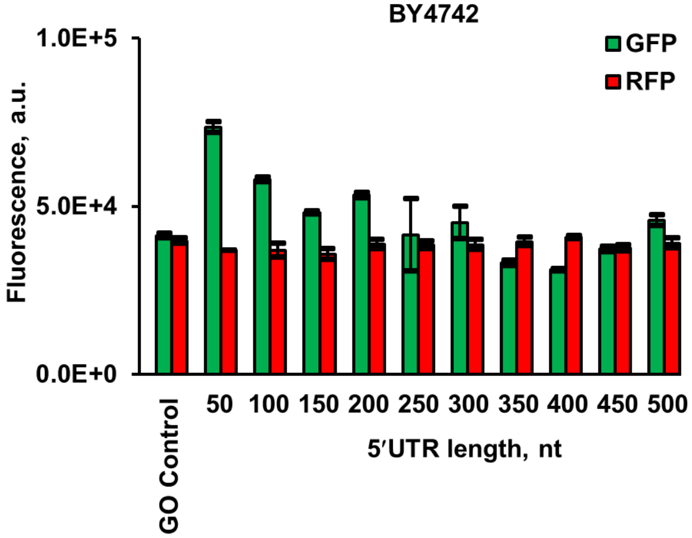


**Figure S3. Mean GFP and RFP fluorescence measured in BY4742 yeast cells, which were transformed with different NUS1 5' UTR RNA-ID constructs as indicated.**

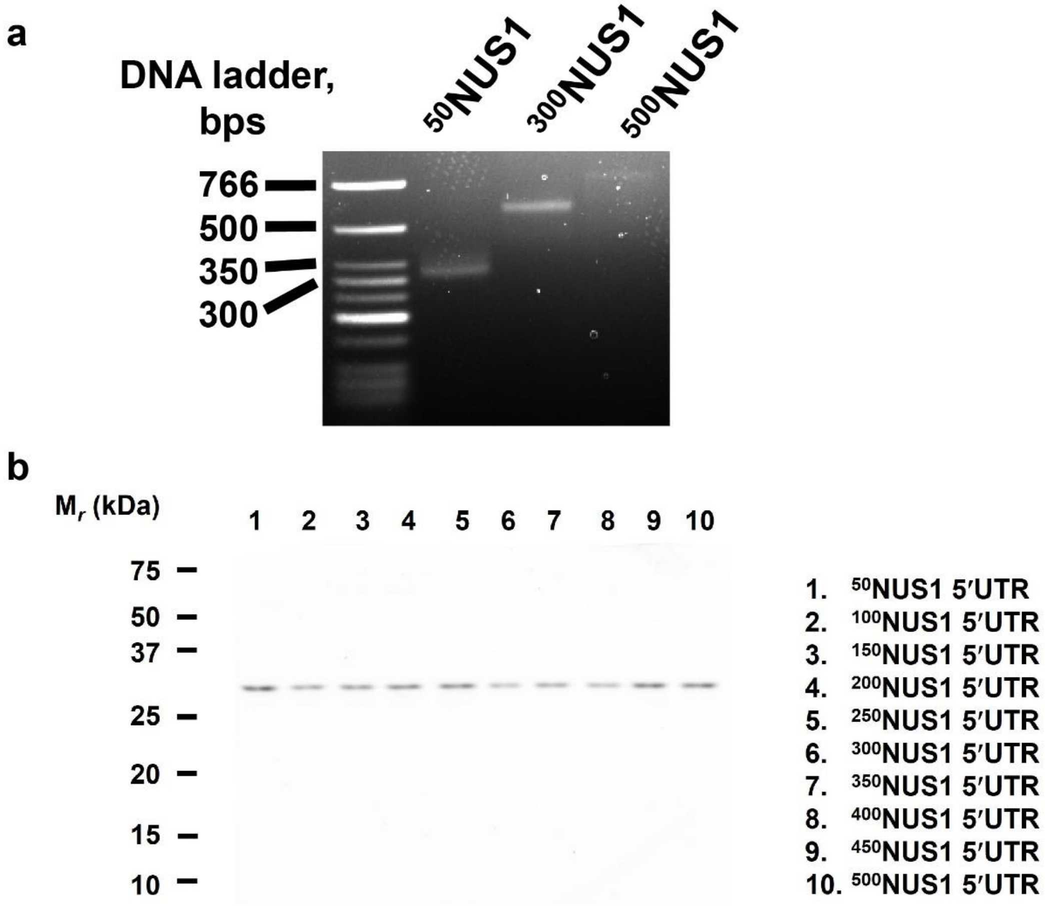


**Figure S4. 5' RACE and SDS-PAGE analysis of different NUS1 5' UTR RNA-ID variants.** (**a**) 5' end of NUS1 5' UTR RNA-ID was analyzed by RACE (Rapid Amplification of cDNA Ends) consisting of sequential dephosphorylation, de-capping, ligation to adaptor RNA (34 nts long) and reverse transcription of yeast total RNA. cDNAs were amplified using a forward primer (HIV-F1) which anneals to adaptor RNA and a reverse primer which anneals to GFP-ORF (5RACE-R2, Supplementary Table S7). PCR products were run on 1% agarose gel. Expected lengths of 50NUS1, 300NUS1 and 500NUS1 amplicons were of 345, 595 and 795 base pairs, respectively. **(b)** Lysates from yeast cells transformed with NUS1 5' UTR RNA-ID variants were run on 15% SDS-PAGE gel and GFP fluorescence was detected by Amersham Typhoon Biomolecule Imager (GE Healthcare, Chicago, IL) with settings of excitation wavelength = 488 nm, emission wavelength = 525 nm, PMT = 400V.

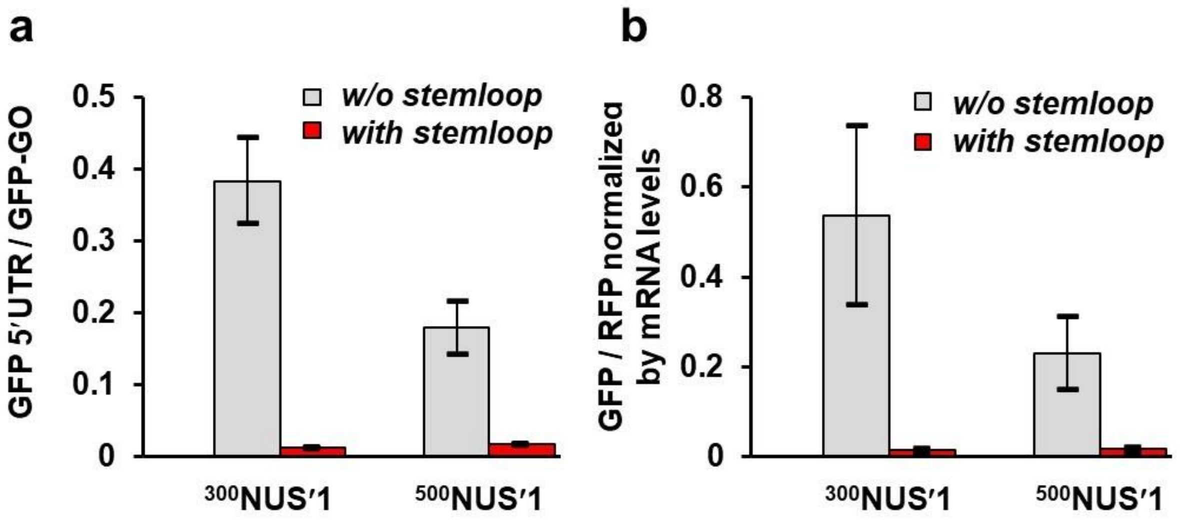


**Figure S5. RNA secondary structure in the 5'UTR strongly inhibits translation of $^{300}$NUS' and $^{500}$NUS' GFP mRNAs.** 54 nucleotides in the middle of $^{300}$NUS' and $^{500}$NUS' 5' UTR were replaced with 44-nt long RNA stem-loop. (**a**) Mean GFP fluorescence was normalized to that in GO RNA-ID reporter lacking both 5' and 3' UTRs. **(b)** Mean GFP fluorescence was normalized to respective mRNA levels, which were determined by RT-qPCR.

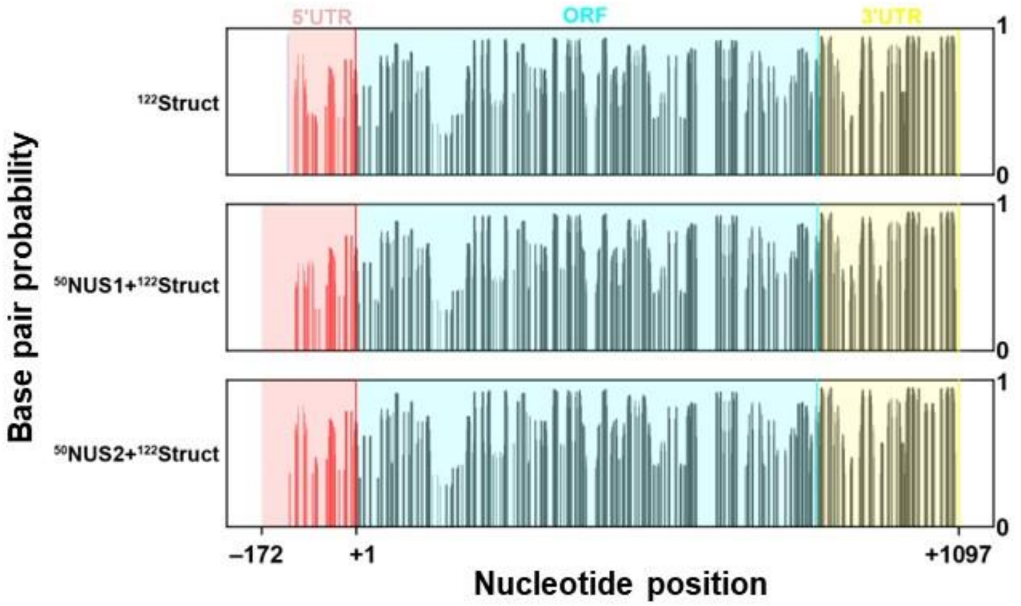


**Figure S6. The nucleotide base pairing probability of ¹²²Struct, NUS3-¹²²Struc, NUS4-¹²²Struct GFP mRNA variants.** The base pairing probabilities were estimated by a partition function calculation. The Y axis is the probability of base pairing. The X axis represents the nucleotide position in the mRNA. The 5′ UTR, ORF and 3′ UTR are highlighted in red, cyan and yellow, respectively.

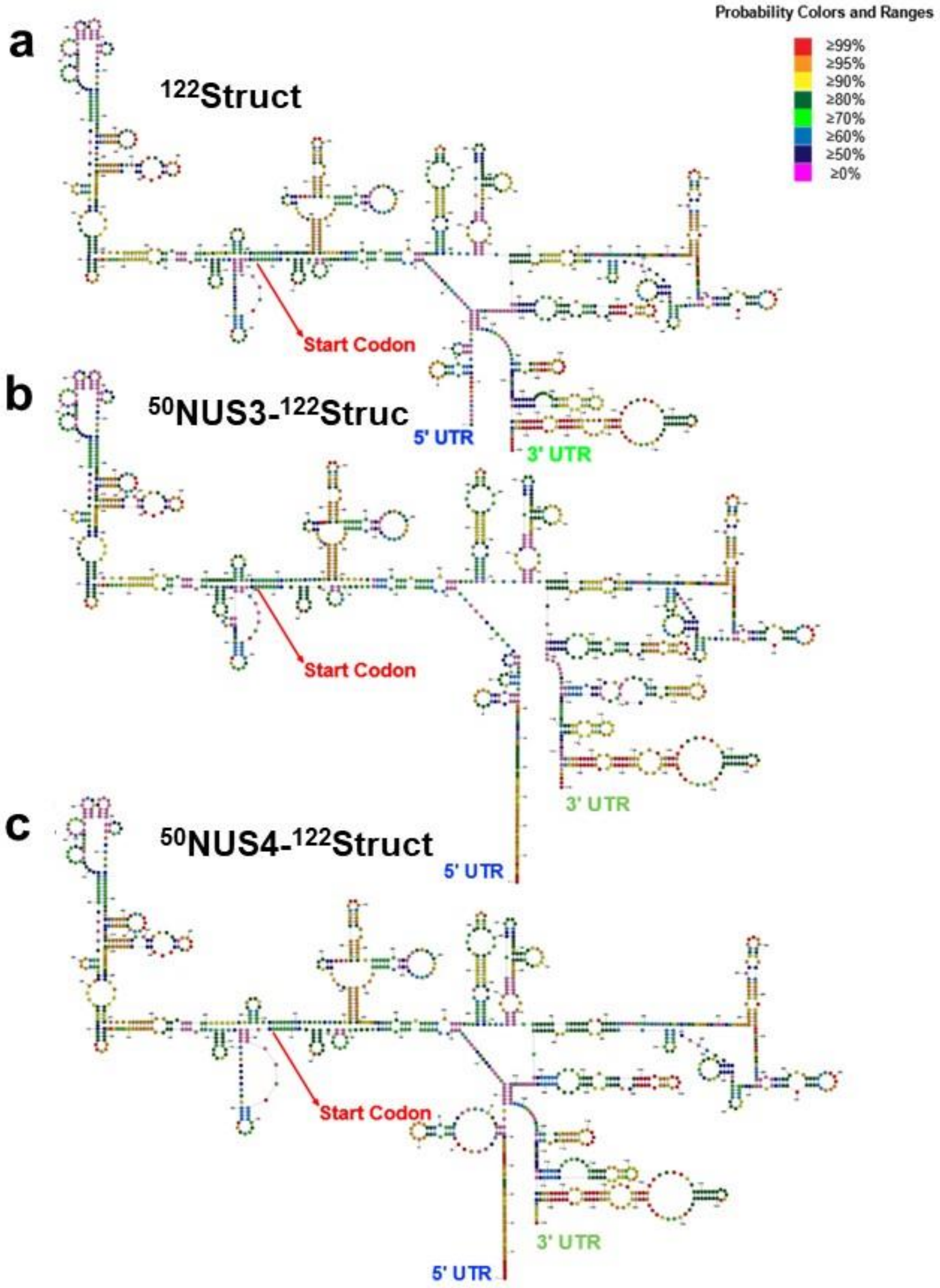


**Figure S7. Secondary structures of $^{122}$Struct, NUS3-$^{122}$Struc, NUS4-$^{122}$Struct GFP mRNA variants.** The predicated lowest free energy structure of $^{122}$Struct (**a**), NUS3-$^{122}$Struc (**b**), NUS4-$^{122}$Struct (**c**) GFP mRNAs. The structures were drawn with *StructureEditor* program from *RNAstructure* software package (https://rna.urmc.rochester.edu/RNAstructure.html). Base pair probabilities, predicated using a partition function, are indicated by the color key.

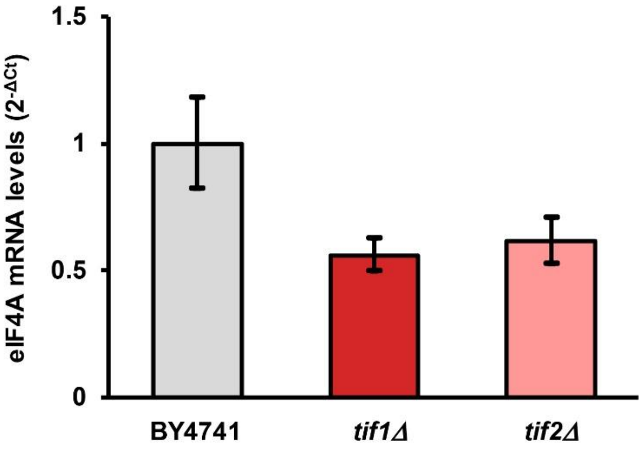


**Figure S8. Changes in eIF4A mRNA levels in *tif1*Δ and *tif2*Δ strains relative to those in wild-type (BY4147) strain, $2^{-\Delta Ct}$, measured by RT-qPCR.**

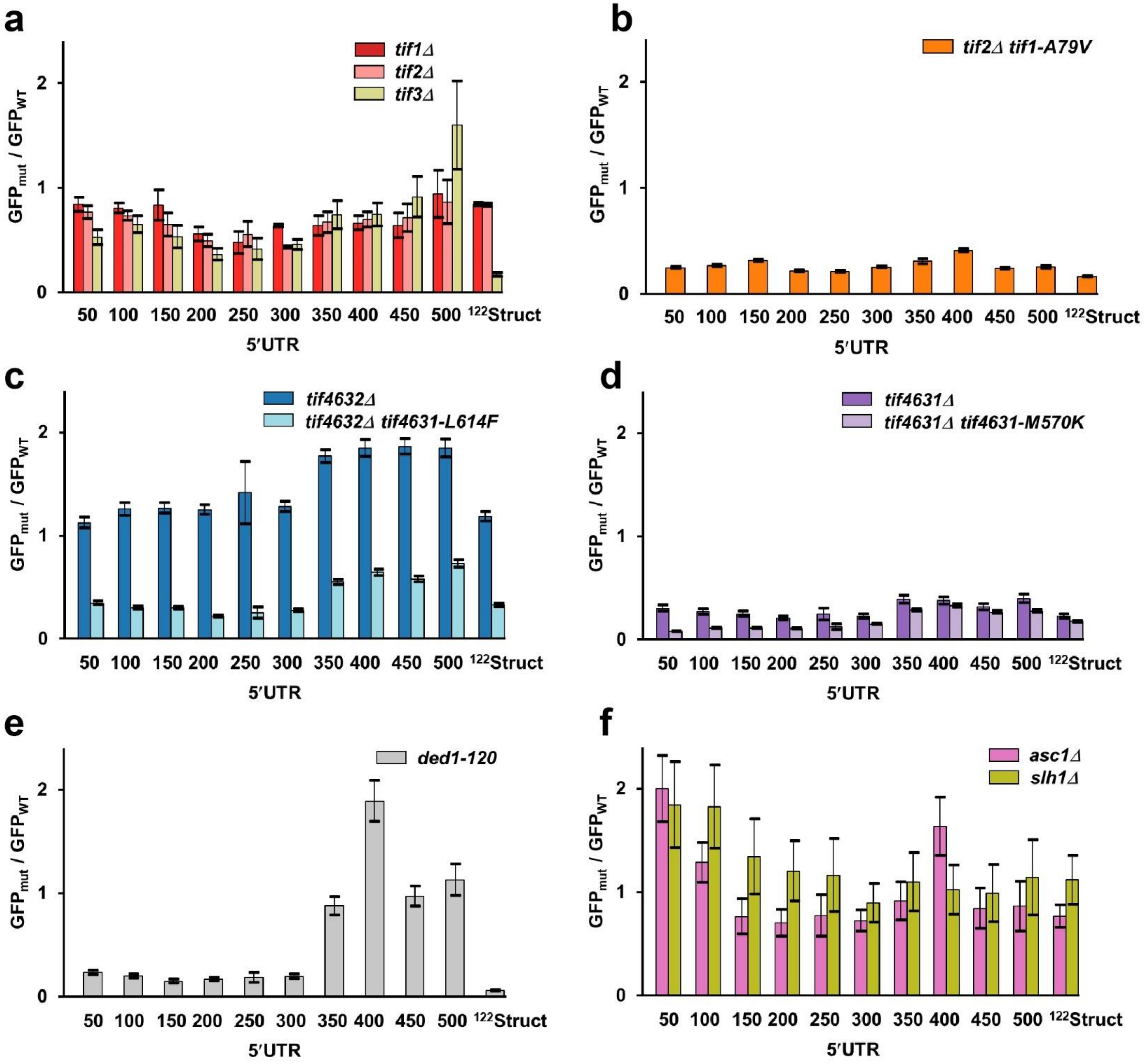


**Figure S9. Mutational perturbations of initiation factors similarly reduce translation of GFP mRNAs with short and long unstructured NUS' 5 'UTRs.** NUS' 5' UTR RNA-ID variants were transformed in yeast strains baring mutations/deletions of initiation factors as indicated. GFP fluorescence measured in mutant strains normalized to respective GFP fluorescence measured in wild-type strain. Error bars indicate standard deviations determined from three biological replicates. (**a**) eIF4A-encoding *tif1, tif2* or eIF4B-encoding *tif3* were deleted in BY4741 background as indicated. **(b)** Either wild-type (*TIF1*) or temperature sensitive (*tif1-A79V*) alleles of eIF4A-encoding *tif1* were expressed in the strain lacking both chromosomal copies of *tif1* and *tif2*. **(c)**

An amino acid substitution L614F, which reduces eIF4G binding to eIF4A, was introduced into *tif4631* in the background of *tif4632* deletion to create temperature-sensitive *tif4632∆tif4631-L614F* strain. **(d)** An amino acid substitution M570K, which reduces eIF4G binding to eIF4A, was introduced into *tif4632* in the background of *tif4631* deletion to create temperature-sensitive *tif4631∆tif4632-M570K* strain. (**e**) Amino acid substitutions G108D and G494D were introduced in Ded1-encoding *ded1* to create *ded1-120* strain. (**f**) In *slh1∆* or *asc1∆* strains, helicase Slh1 and ribosomal protein Asc1, respectively.

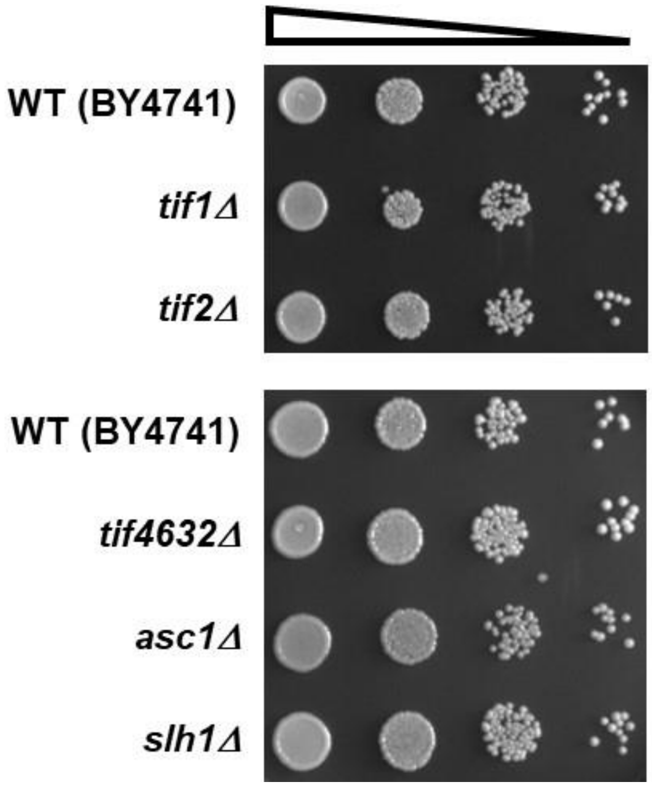


**Figure S10. Deletion of *tif1, tif2, tif4632, asc1* or *slh1* does not affect cell growth on YPD plates.** Strains were growth on YPD media for 48 hours at 30°C.

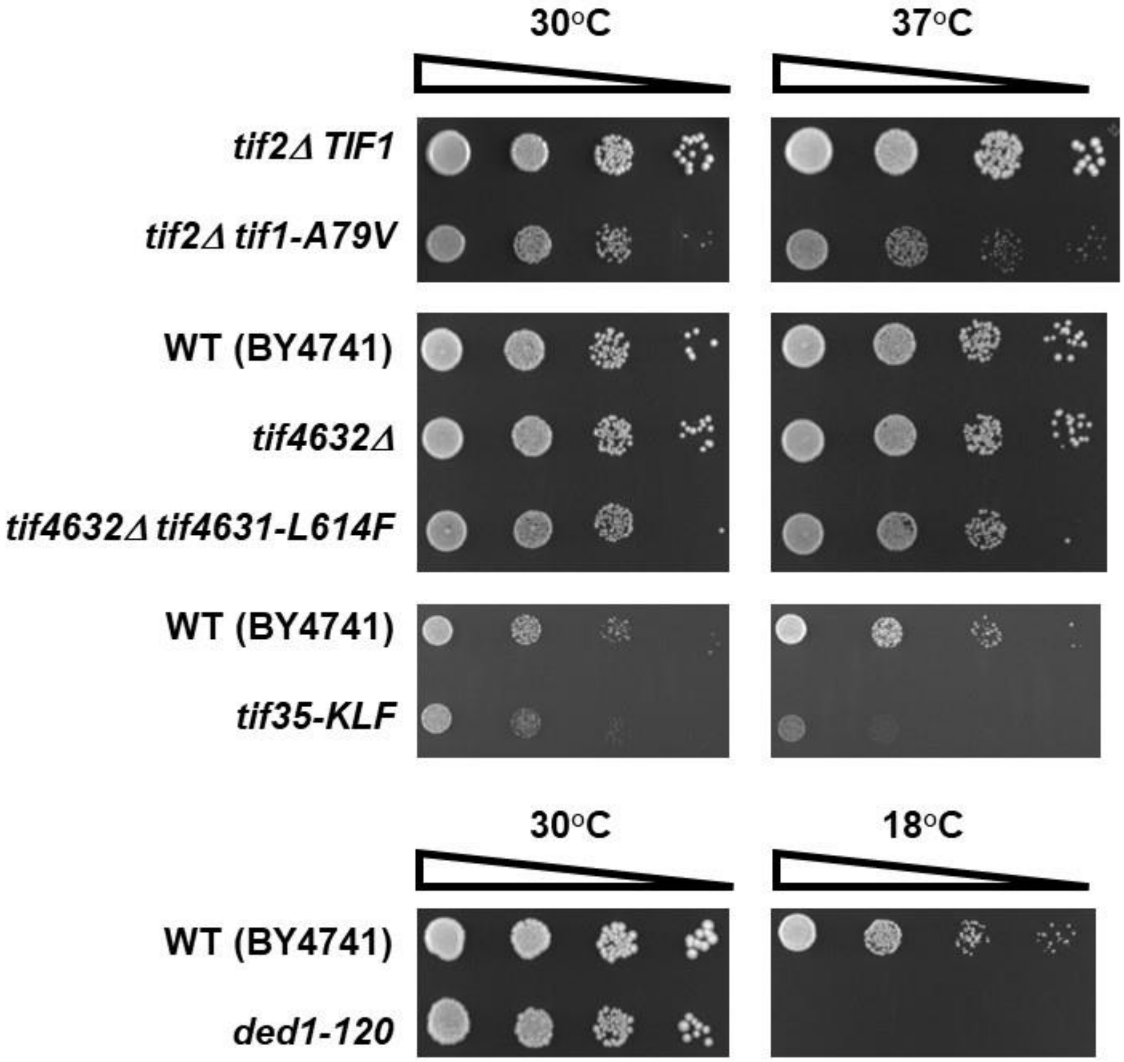


**Figure S11. Temperature and cold-sensitive mutations in *tif1*, *tif4631*, *tif35* and *ded1* slowed yeast growth at non-permissive temperatures.** Mutant strains were growth for 48 hours on YPD media at 30, 37 or 18°C as indicated.

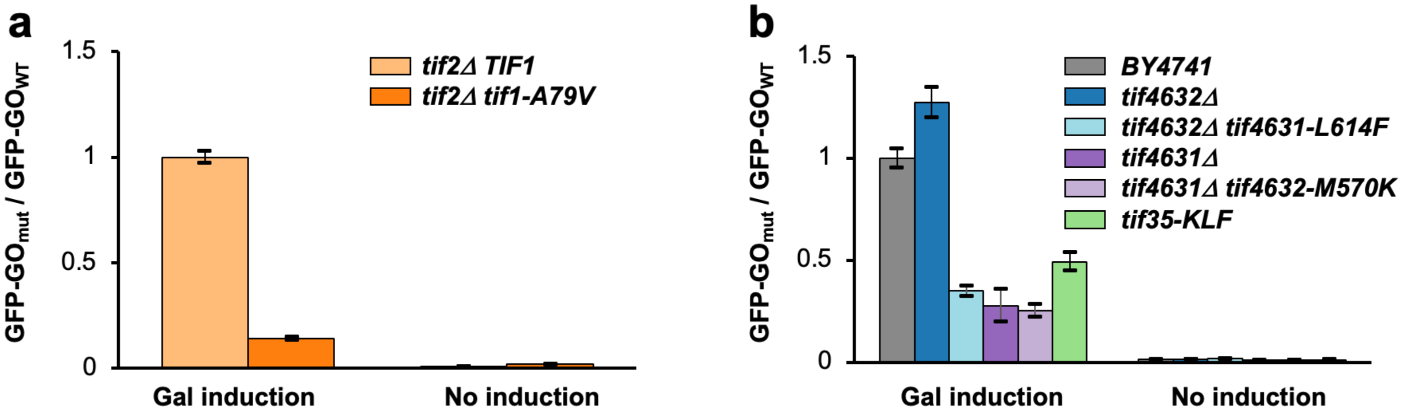


**Figure S12. GFP is not synthesized in the absence of galactose.** GFP-GO RNA-ID construct was transformed in yeast strains barring temperature sensitive mutations or deletions of initiation factors as indicated. GFP fluorescence measured in mutant strains normalized to respective GFP fluorescence measured in wild-type (BY4147 or *tif2*Δ*TIF1*) strain. Galactose was added to induce *Gal1,10* promoter concurrently to switching from 30 to 37°C. Error bars indicate standard deviations determined from three biological replicates.

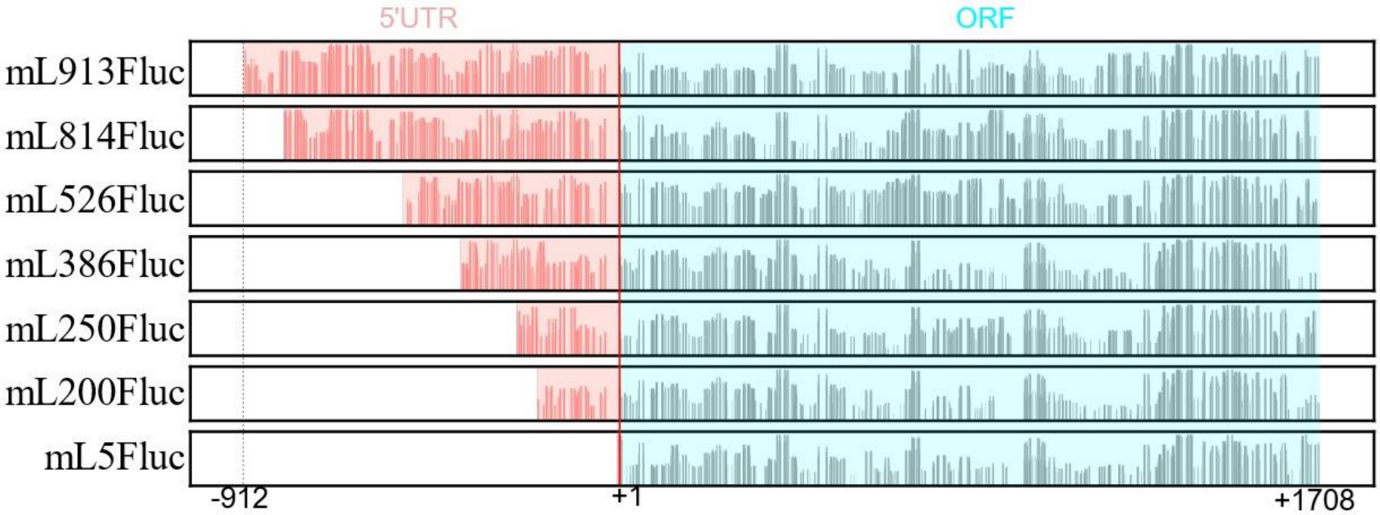


**Figure S13. The nucleotide base pairing probability of reporter mRNAs used by Vassilenko et al. (1).** The base pairing probabilities were estimated with a partition function calculation. The Y axis is the probability of base pairing. The X axis represents the nucleotide position in the mRNA. The 5′ UTR and ORF are highlighted in red and cyan, respectively.

#### Supplementary Methods

##### Yeast genomic DNA (gDNA) purification.

Cultured yeast cells on YPD plate were collected and lysed by vortexing for 5 min in 10 mM Tris-HCl, pH 8.0, 1 mM EDTA, 2% Triton-X 100, 1% SDS, 100 mM NaCl with micro-glass beads. The supernatant was collected, and 8-fold excess volume of 100% ethanol was added and incubated for 1 hr at -20°C. After centrifuge at 1330 rpm for 10 min, supernatant was removed. The precipitate was dissolved in 400 µl 10 mM Tris-HCl, pH 8.0, 1 mM EDTA with 1 µl RNase A (10 mg/mL) and incubated for 5 min at 23°C. The pure genomic DNA was collected by ethanol precipitation.

##### Cloning of RNA-ID plasmid containing desired sequences at 5′UTR of GFP-ORF.

The original RNA-ID plasmid contains *MET15* sequence for auxotrophic selection (2). Since we intended to use different yeast strains (BY4741 and BY4742) and BY4742 contains native *MET15* gene, we modified the original RNA-ID plasmid by swapping the *MET15* sequence with the *URA3* sequence by Gibson assembly. *MET15* eliminated RNA-ID DNA was amplified by PCR using original RNA-ID plasmid as a template DNA and primers (*URA3*-Replace-Vec-F1 and *URA3*-Replace-Vec-R1). Insert *URA3* DNAs were amplified using S288C gDNA as a template DNA. At the same time, we eliminated the StuI site present in native *URA3* gene with synonymous mutation codon by doing two separate PCR reactions. Upstream of *URA3* gene was amplified using *URA3*-replace-ins-R1 and StuI-remove-R1 primers. Downstream of *URA3* gene was amplified using *URA3*-replace-ins-F1 and StuI-remove-F1 primers. *MET15* eliminated linear RNA-ID DNA, upstream of *URA3*, and downstream of *URA3* were combined by Gibson assembly and obtained RNA-ID-*URA3* plasmid. For further cloning we used this RNA-ID plasmid.

$^{50}$NUS 5′UTR ~ $^{500}$NUS 5′UTR containing RNA-ID-plasmid construction was done as follows. We first constructed the RNA-ID plasmid carrying $^{500}$NUS 5′UTR. The $^{500}$NUS 5′UTR DNA fragment containing 15 bp 3' end of *Gal1,10* promoter at the 5' end and 31 bp 5' end of tagged GFP-ORF sequence at the 3' end was produced by PCR reaction combining 3 synthesized DNA oligo nucleotides with a size of 200 bp in the presence of forward primer (NR-5UTR-F1) and reverse primers (500NR-5UTR-R1 and 500NR-5UTR-R2). Also upstream flanking DNA fragment and downstream flanking DNA fragment were produced by PCR. Then the upstream DNA, $^{500}$NUS 5′UTR DNA, and downstream DNA were combined by Gibson assembly. Assembled DNA was further amplified by PCR using SpeI-F1 and SalI-R1 primers, agarose-gel purified, and digested by SpeI/SalI ($^{500}$NUS 5′UTR insert DNA). The original RNA-ID plasmid was also digested by SpeI/SalI and vector part of DNA was agarose-gel purified (RNA-ID vector). Finally, the complete $^{500}$NUS 5′UTR-RNA-ID plasmid was generated by ligation of $^{500}$NUS 5′UTR insert DNA and RNA-ID vector. Obtained plasmid was verified by Sanger sequencing (Genewiz-Azenta). Then we generated the rest of plasmid ($^{50}$NUS 5′UTR-RNA-ID ~ $^{450}$NUS 5′UTR-RNA-ID) as follows. Upstream and downstream insert DNA fragments were produced by PCR using $^{500}$NUS 5′UTR-RNA-ID plasmid as a template DNA and primer sets (SpeI-F1/50NR-5UTR-R1 ~ 450NR-5UTR-R1 for upstream DNA and 50NR-5UTR-F1 ~ 450NR-5UTR-F1/SalI-R1 for upstream DNA). These upstream and downstream DNAs were combined by Gibson assembly, digested by SpeI/SalI, and ligated with SpeI/SalI digested vector part of RNA-ID DNA. Obtained plasmid was verified by Sanger sequencing (Genewiz-Azenta).

The $^{300}$NUS-SL 5′UTR containing the RNA-ID-plasmid construction was done as follows. Using $^{300}$NUS RNA-ID plasmid as a template DNA, we performed PCR to amplify an upstream of insert DNA fragment with primers (SpeI-F1, 50SL2-300NR-5UTR-DNA1-R1, and 50SL2-300NR-5UTR-DNA1-R2), and a downstream of insert DNA fragment with primers (Sal1-R1, 50SL2-300NR-5UTR-DNA1-F1, and 50SL2-300NR-5UTR-DNA1-F2). The upstream of insert DNA fragment was digested by SpeI/AflII and the downstream of insert DNA fragment was digested by SalI/AflII. Finally, the digested upstream of insert DNA fragment, the downstream of insert DNA fragment, and the RNAID vector DNA (see above) were ligated to create the $^{300}$NUS-SL 5′UTR RNA-ID plasmid. Obtained plasmid was verified by Sanger sequencing (Genewiz-Azenta).

$^{50}$NUS3-$^{122}$Struct 5′UTR and $^{50}$NUS4-$^{122}$Struct containing the RNA-ID-plasmid construction was done as follows. Upstream and downstream insert DNA fragments were produced by PCR using the $^{122}$Struct 5′UTR-RNA-ID plasmid as a template DNA. For producing $^{50}$NUS3-$^{122}$Struct 5′UTR insert DNAs, SpeI-F1 and RNAID-Vec-5'-350-R1 were used to produce the upstream part of the insert DNA. Also, RNAID-Vec-5'-GAPDH-50-F1, GAPDH-50-F2, and SalI-R1 were used to produce the downstream part of insert DNA. For producing $^{50}$NUS4-$^{122}$Struct 5′UTR insert DNAs, SpeI-F1 and GAPDH-50RE-R2 were used to produce upstream part of the insert DNA. Also, GAPDH-50RE-F1, GAPDH-50RE-F2, and SalI-R1 were used to produce the downstream part of insert DNA. Then, these upstream and downstream DNAs were combined by Gibson assembly, digested by SpeI/SalI, and ligated with the SpeI/SalI-digested vector part of RNA-ID DNA. Obtained plasmid was verified by Sanger sequencing (Genewiz-Azenta).

**Cloning of plasmid for gene mutation.**

At first, DNA fragments for Gibson assembly were produced by PCR using the primer-DNA template pairs as listed in **Supplementary Tables 2-12**. Next, appropriate DNA segment sets and corresponding plasmid vector were combined by Gibson assembly to generate the complete plasmid. Obtained plasmid was verified by Sanger sequencing (Genewiz-Azenta). Purified plasmids were digested by restriction enzymes as indicated in **Supplementary Table 13** and DNA fragments other than the carrier vector part were agarose-gel purified.

**SDS-PAGE analysis with fluorescence detection.**

Cells were lysed in 1.5 mL tubes containing 200 µL of 50 mM HEPES, pH 7.5, 1 mM EDTA, 0.5% Triton X-100, 1 mM DTT, 10% glycerol, 1 M NaCl, Thermo Scientific Pierce Protease Inhibitor Mini Tablets, EDTA-free (1 tablet in 50 mL, Fisher Scientific), and 1 mM PMSF (3). The solution was mixed with 300 mg acid-washed Glass Beads (Sigma-Aldrich, St. Louis, MO) and vortexed for 10 min at 4°C. After centrifuging at 13,300 rpm for 10 min, supernatant was collected. ~20 $Abs_{260}$ of lysate were mixed with equal volume of 2X Laemmli buffer containing beta-mercaptoethanol and loaded onto the 15% SDS-PAGE gel. After running at 150V for 100 min, the gel was visualized by Amersham Typhoon Biomolecule Imager (GE Healthcare, Chicago, IL) with excitation at 488 nm, emission filter set to 525 nm and PMT set to 400V.

**RT-qPCR.**

RT-qPCR was performed according to the method described previously (4) using primers provided in **Supplementary Table 14**. Yeast total RNA was prepared by using formamide-EDTA as described elsewhere (5). cDNA was synthetized using Superscript II Reverse Transcriptase (Invitrogen). cDNA was amplified using Fast SYBR Green Master Mix (Applied Biosystems), detected using the 7500 Fast Real-Time PCR system, and analyzed with the 7500 Software v2.3 (Applied Biosystems).

To validate TIF1 and TIF2 gene transcription, we used TFC1, TAF10, and ALG9 as reference genes (6). Primer pair used to quantify TIF1 or TIF2 transcripts was tif1-2-semiqPCR-F1/ tif1-2-qPCR-R1 and reference primer pairs were TFC1-F1/TFC-R1, TAF10-F1/TAF10-R1, and ALG9-F1/ALG10-R1. Primer pairs used to quantify GFP transcript and reference RFP transcripts were GFP-qPCR-F4/GFP-qPCR-R4 and RFP-qPCR-F5/RFP-qPCR-R5, respectively.

**5′RACE.**

Approximately 10 µg of yeast total RNA from $^{50}$NUS, $^{300}$NUS, or $^{500}$NUS RNA-ID DNA transformed BY4741 cells were first treated with Calf Intestinal Alkaline Phosphatase (NEB) to dephosphorylate RNAs without cap. Then the cap on the mRNA was removed by mRNA Decapping Enzyme (NEB), leaving phosphate bound at 5' end. Adaptor RNA was synthesized by IDT. The RNA sequence is 5′-UUCUUCUGAAGAUAAAGCAACAACAACAAGGCAA -3′. Ligation of adaptor RNA to the samples was performed using T4 RNA ligase I (NEB). Reverse transcription was performed by the method (see RT-qPCR section) and each target cDNA was amplified using a forward primer (HIV-F1) that anneals to adaptor RNA and a reverse primer that anneals to GFP-ORF (5RACE-R2, **Supplementary Table 14**). Since some transcription starts not exactly at the expected 5' end of the 5'UTR, we cloned the 5'RACE amplicon into an pSP64 plasmid to validate sequence by Sanger sequencing. In order to clone the 5'RACE amplicon into pSP64 using SacI and HindIII site, we created a SacI site at 5' end of the 5'RACE amplicon by PCR using a primer set (NR-Lig-SacI-F2-2, NR-Lig-SacI-F3, and 5RACE-R2). There is a HindIII site ~50bp downstream from the 3' end of 5' UTR. After PCR, SacI/HindIII restriction digestion, each purified DNA was ligated with SacI/HindIII digested pSP64 plasmid. We collected ~10 ligated plasmid clones for each $^{50}$NUS, $^{300}$NUS, or $^{500}$NUS RNA-ID sample and Sanger sequencing was performed at Genewize-Azenta using a pSP64 specific primer (M13R-Hiro).

**Spot assay.**

A small volume (5 ml) Yeast cells were cultured in YPD medium overnight at 30°C, adjusted concentration at $OD_{600}$~0.2, and cultured 4 hrs in YPD medium at indicated temperature. Cells were appropriately diluted in YPD and 5 µl of each dilution was spotted on YPD solid Medium. After indicated time (1 – 4 days) plates were photographed with a Molecular Imager Gel Doc XR+ system (Bio-Rad, Hercules, CA).

**Supplementary Table 1. 5' UTR sequences of GFP mRNA in RNA-ID constructs.**

| 5' UTR | Sequence (5' to 3') | Linguistic complexity† |
|---|---|---|
| GO | GCUAGC | 0.600 |
| 50NUS1 | CACCAAAAAACACACACACAACAUAAAACACAAAAACAAAACAACACC UAAAA | 0.0278 |
| 100NUS1 | CACCAAAAAACACACACACAACAUAAAACACAAAAACAAAACAACACC UAAACAACACACACACACACACACACACACACACACACAAAACUAC ACAAAAA | 0.00494 |
| 150NUS1 | CACCAAAAAACACACACACAACAUAAAACACAAAAACAAAACAACACC UAAACAACACACACACACACACACACACACACACACACAAAACUAC ACAACAAACAACAACAAAACACAAAACACACACAACACACACACAACA CAACAAAAA | 0.00226 |
| 200NUS1 | CACCAAAAAACACACACACAACAUAAAACACAAAAACAAAACAACACC UAAACAACACACACACACACACACACACACACACACACAAAACUAC ACAACAAACAACAACAAAACACAAAACACACACAACACACACACAACA CAACAACAAAAAACACAUACUAAACACAACACAAAAAACACACAACAC ACAACACAAAA | 0.00160 |
| 250NUS1 | CACCAAAAAACACACACACAACAUAAAACACAAAAACAAAACAACACC UAAACAACACACACACACACACACACACACACACACACAAAACUAC ACAACAAACAACAACAAAACACAAAACACACACAACACACACACAACA CAACAACAAAAAACACAUACUAAACACAACACAAAAAACACACAACAC ACAACACACAACAACAAACCCCCACACCCCCCCCCUACAAAAACACA AAAAACACACAAA | 0.00182 |
| 300NUS1 | CACCAAAAAACACACACACAACAUAAAACACAAAAACAAAACAACACC UAAACAACACACACACACACACACACACACACACACACAAAACUAC ACAACAAACAACAACAAAACACAAAACACACACAACACACACACAACA CAACAACAAAAAACACAUACUAAACACAACACAAAAAACACACAACAC ACAACACACAACAACAAACCCCCACACCCCCCCCCUACAAAAACACA AAAAACACACACACUACACACAUCAUUACAACAAUCACAACCUCACAC AACCCUAUACCAAAA | 0.00216 |
| 350NUS1 | CACCAAAAAACACACACACAACAUAAAACACAAAAACAAAACAACACC UAAACAACACACACACACACACACACACACACACACACAAAACUAC ACAACAAACAACAACAAAACACAAAACACACACAACACACACACAACA CAACAACAAAAAACACAUACUAAACACAACACAAAAAACACACAACAC ACAACACACAACAACAAACCCCCACACCCCCCCCCUACAAAAACACA AAAAACACACACACUACACACAUCAUUACAACAAUCACAACCUCACAC AACCCUAUACCAAAACAAAAAACCCCUACACAAAAAACACACACACAA CUACACUAAAUCUCAAA | 0.00221 |
| 400NUS1 | CACCAAAAAACACACACACAACAUAAAACACAAAAACAAAACAACACC UAAACAACACACACACACACACACACACACACACACACAAAACUAC ACAACAAACAACAACAAAACACAAAACACACACAACACACACACAACA CAACAACAAAAAACACAUACUAAACACAACACAAAAAACACACAACAC ACAACACACAACAACAAACCCCCACACCCCCCCCCUACAAAAACACA AAAAACACACACACUACACACAUCAUUACAACAAUCACAACCUCACAC AACCCUAUACCAAAACAAAAAACCCCUACACAAAAAACACACACACAA CUACACUAAAUCUCACAAACAACAACACAAAACACAACCUACACACAC AAAACACAACCCCUCAAAA | 0.00182 |
| 450NUS1 | CACCAAAAAACACACACACAACAUAAAACACAAAAACAAAACAACACC UAAACAACACACACACACACACACACACACACACACACAAAACUAC ACAACAAACAACAACAAAACACAAAACACACACAACACACACACAACA CAACAACAAAAAACACAUACUAAACACAACACAAAAAACACACAACAC ACAACACACAACAACAAACCCCCACACCCCCCCCCUACAAAAACACA | 0.00186 |

| | | |
|---|---|---|
| | AAAAACACACACACUACACACAUCAUUACAACAAUCACAACCUCACAC AACCCUAUACCAAAACAAAAAACCCCUACACAAAAAACACACACACAA CUACACUAAAUCUCACAAACAACAACACAAAACACAACCUACACACAC AAAACACAACCCCUCACUAUUAUACACACACAAAAAAUACUAAACAAA AACAAAAAAUAAAAAACAAAA | |
| $^{500}$NUS1 | CACCAAAAAACACACACACAACAUAAAACACAAAAACAAAACAACACC UAAACAACACACACACACACACACACACACACACACACACAAAACUAC ACAACAAACAACAACAAAACACAAAACACACACAACACACACACAACA CAACAACAAAAAACACAUACUAAACACAACACAAAAAACACACAACAC ACAACACACAACAACAAACCCCCACACCCCCCCCCUACAAAAACACA AAAAACACACACACUACACACAUCAUUACAACAAUCACAACCUCACAC AACCCUAUACCAAAACAAAAAACCCCUACACAAAAAACACACACACAA CUACACUAAAUCUCACAAACAACAACACAAAACACAACCUACACACAC AAAACACAACCCCUCACUAUUAUACACACACAAAAAAUACUAAACAAA AACAAAAAAUAAAAAACACACAAAAACCCCCUACUAAAACAAAAACA AAAAUAAAUAAAAAAUACCCAAA | 0.00168 |
| $^{300}$NUS2 | ACACACUACACACACCAAACCCAUAAUCACAACACACACAAUUAAAUA CCAACUCUAAACACAAAAACUAUCAUUACACACACCAAAUUUCUCAUA CUACUACUCACUAUUACACAAACACCUCAUUUAUAACAACAACAUUC AUUUCUACAAAACACUCAUCAAACCUUACACAUACAACCCCAAUUAC CUUUUUACCAAUCACAAUACACAAAACCUACUAAACCCCCCAACCCU ACAACAUCUCAUACACACACAAAUACAAACAACACACUUAAACAACAA CAACCCCUAAAACCUAAA | 0.0150 |
| $^{300}$NUS1 +SL | ACACACUACACACACCAAACCCAUAAUCACAACACACACAAUUAAAUA CCAACUCUAAACACAAAAACUAUCAUUACACACACCAAAUUUCUCAUA CUACUACUCACUAUUACACAAACACCUCAUUUAUAACAACAACAUUC AUUUCUAAGACGCUUGCUUGUCACCGCUCCUUAAGGAGUGGUGGCA GGUAGGCGAAAAUCACAAUACACAAAACCUACUAAACCCCCCAACCC UACAACAUCUCAUACACACACAAAUACAAACAACACACUUAAACAACA ACAACCCCUAAAACCUAAA | 0.144 |
| $^{122}$Struct | GCUAGCCCCCCGGUUUCUAUAAAUUGAGCCCGCAGCCUCCCGCUUC GCUCUCUGCUCCUCCUGUUCGACAGUCAGCCGCAUCUUCUUUUGC GUCGCCAGCCGAGCCACAUCGCUCAGACACC | 0.441 |
| $^{50}$NUS3 - $^{122}$Struct | CAAUAAAAACCUCCCUAAAACCCCAAUACUCCCUACUAAUAAAACCC CCCGCUAGCCCCCCGGUUUCUAUAAAUUGAGCCCGCAGCCUCCCGC UUCGCUCUCUGCUCCUCCUGUUCGACAGUCAGCCGCAUCUUCUUUU GCGUCGCCAGCCGAGCCACAUCGCUCAGACACC | 0.373 |
| $^{50}$NUS4 - $^{122}$Struct | CACCCCCUAAAUAAUCACAACCACACUCACAUUUCCUACCCCCCCCA UAAGCUAGCCCCCCGGUUUCUAUAAAUUGAGCCCGCAGCCUCCCGC UUCGCUCUCUGCUCCUCCUGUUCGACAGUCAGCCGCAUCUUCUUUU GCGUCGCCAGCCGAGCCACAUCGCUCAGACACC | 0.452 |
| $^{50}$NUS′ | CCCAACACUUACCCCCCCUACUCCCCAAAACCCCAUCCAAUACUCCC AAC | 0.112 |
| $^{100}$NUS′ | AACUACCCCAACAAAAACAAACACUACCCCCCAAUCUCUAUCCCCCA CCCCCCAACACUUACCCCCCCUACUCCCCAAAACCCCAUCCAAUACU CCCAACAAA | 0.0351 |
| $^{150}$NUS′ | CCCUAAAACAAAACCCCCCAACCAACAAUCCUCAACCUACCAAAACAA ACAACUACCCCAACAAAAACAAACACUACCCCCCAAUCUCUAUCCCC CACCCCCCAACACUUACCCCCCCUACUCCCCAAAACCCCAUCCAAUA CUCCCAACAAA | 0.0265 |
| $^{200}$NUS′ | ACCCCCAACCAACAAAACACAUAUAACACUCCCUCACAAAUUCCCCC ACACCCUAAAACAAAACCCCCCAACCAACAAUCCUCAACCUACCAAAA CAAACAACUACCCCAACAAAAACAAACACUACCCCCCAAUCUCUAUC | 0.0262 |

| | | |
|---|---|---|
| | CCCCACCCCCCAACACUUACCCCCCCUACUCCCCAAAACCCCAUCCA<br>AUACUCCCAACAAA | |
| $^{250}$NUS′ | AACCCAACAACAACAAACAAAAACUCACAACCAAAUCUCACACACAAC<br>CAACCCCCAACCAACAAAACACAUAUAACACUCCCUCACAAAUUCCC<br>CCACACCCUAAAACAAAACCCCCCAACCAACAAUCCUCAACCUACCA<br>AAACAAACAACUACCCCAACAAAAACAAACACUACCCCCCAAUCUCUA<br>UCCCCCACCCCCCAACACUUACCCCCCCUACUCCCCAAAACCCCAUC<br>CAAUACUCCCAACAAA | 0.0178 |
| $^{300}$NUS′ | ACAUACCCCACACUCAACCCAUACCUAAAACCCCCACUCAACUCAAA<br>ACCAACCCAACAACAACAAACAAAAACUCACAACCAAAUCUCACACAC<br>AACCAACCCCCAACCAACAAAACACAUAUAACACUCCCUCACAAAUUC<br>CCCCACACCCUAAAACAAAACCCCCCAACCAACAAUCCUCAACCUAC<br>CAAAACAAACAACUACCCCAACAAAAACAAACACUACCCCCCAAUCUC<br>UAUCCCCCACCCCCCAACACUUACCCCCCCUACUCCCCAAAACCCCA<br>UCCAAUACUCCCAACAAA | 0.00861 |
| $^{350}$NUS′ | CAAUAAAAACCUCCCUAAAACCCCAAUACUCCCUACUAAUAAAACCC<br>CCCACAUACCCCACACUCAACCCAUACCUAAAACCCCCACUCAACUC<br>AAAACCAACCCAACAACAACAAACAAAAACUCACAACCAAAUCUCACA<br>CACAACCAACCCCCAACCAACAAAACACAUAUAACACUCCCUCACAAA<br>UUCCCCCACACCCUAAAACAAAACCCCCCAACCAACAAUCCUCAACC<br>UACCAAAACAAACAACUACCCCAACAAAAACAAACACUACCCCCCAAU<br>CUCUAUCCCCCACCCCCCAACACUUACCCCCCCUACUCCCCAAAACC<br>CCAUCCAAUACUCCCAACAAA | 0.00750 |
| $^{400}$NUS′ | ACAAUACUCCUACAACACAAACAAAAUAAAACACUCACACAAAACAAC<br>UACAAUAAAAACCUCCCUAAAACCCCAAUACUCCCUACUAAUAAAACC<br>CCCCACAUACCCCACACUCAACCCAUACCUAAAACCCCCACUCAACU<br>CAAAACCAACCCAACAACAACAAACAAAAACUCACAACCAAAUCUCAC<br>ACACAACCAACCCCCAACCAACAAAACACAUAUAACACUCCCUCACAA<br>AUUCCCCCACACCCUAAAACAAAACCCCCCAACCAACAAUCCUCAAC<br>CUACCAAAACAAACAACUACCCCAACAAAAACAAACACUACCCCCCAA<br>UCUCUAUCCCCCACCCCCCAACACUUACCCCCCCUACUCCCCAAAAC<br>CCCAUCCAAUACUCCCAACAAA | 0.00652 |
| $^{450}$NUS′ | CCCACACUAACUACUCACAACCACACUCACAUUUCCUACCCCACACA<br>CACACAAUACUCCUACAACACAAACAAAAUAAAACACUCACACAAAAC<br>AACUACAAUAAAAACCUCCCUAAAACCCCAAUACUCCCUACUAAUAAA<br>ACCCCCCACAUACCCCACACUCAACCCAUACCUAAAACCCCCACUCA<br>ACUCAAAACCAACCCAACAACAACAAACAAAAACUCACAACCAAAUCU<br>CACACACAACCAACCCCCAACCAACAAAACACAUAUAACACUCCCUCA<br>CAAAUUCCCCCACACCCUAAAACAAAACCCCCCAACCAACAAUCCUC<br>AACCUACCAAAACAAACAACUACCCCAACAAAAACAAACACUACCCCC<br>CAAUCUCUAUCCCCCACCCCCCAACACUUACCCCCCCUACUCCCCAA<br>AACCCCAUCCAAUACUCCCAACAAA | 0.00645 |
| $^{500}$NUS′ | ACAACUAAAACCCACCAAUAACUAAAACAACCACAACAUAUCACUCCU<br>ACCCCACACUAACUACUCACAACCACACUCACAUUUCCUACCCCACA<br>CACACACAAUACUCCUACAACACAAACAAAAUAAAACACUCACACAAA<br>ACAACUACAAUAAAAACCUCCCUAAAACCCCAAUACUCCCUACUAAUA<br>AAACCCCCCACAUACCCCACACUCAACCCAUACCUAAAACCCCCACU<br>CAACUCAAAACCAACCCAACAACAACAAACAAAAACUCACAACCAAAU<br>CUCACACACAACCAACCCCCAACCAACAAAACACAUAUAACACUCCC<br>UCACAAAUUCCCCCACACCCUAAAACAAAACCCCCCAACCAACAAUC<br>CUCAACCUACCAAAACAAACAACUACCCCAACAAAAACAAACACUACC<br>CCCCAAUCUCUAUCCCCCACCCCCCAACACUUACCCCCCCUACUCCC<br>CAAAACCCCAUCCAAUACUCCCAACAAA | 0.00596 |

| | | |
|---|---|---|
| $^{300}$NUS′<br>+SL | ACAUACCCCACACUCAACCCAUACCUAAAACCCCCACUCAACUCAAA<br>ACCAACCCAACAACAACAAACAAAAACUCACAACCAAAUCUCACACAC<br>AACCAACCCCCAACCAACAAAACACAUAUAACACUCCCUCACAAAUUC<br>CCCCACAAGACGCUUGCUUGUCACCGCUCCUUAAGGAGUGGUGGCA<br>GGUAGGCGAAAAACUACCCCAACAAAAACAAACACUACCCCCCAAUC<br>UCUAUCCCCCACCCCCCAACACUUACCCCCCCUACUCCCCAAAACCC<br>CAUCCAAUACUCCCAACAAA | 0.110 |
| $^{500}$NUS′<br>+SL | ACAACUAAAACCCACCAAUAACUAAAACAACCACAACAUAUCACUCCU<br>ACCCCACACUAACUACUCACAACCACACUCACAUUUCCUACCCCACA<br>CACACACAAUACUCCUACAACACAAACAAAAUAAAACACUCACACAAA<br>ACAACUAAGACGCUUGCUUGUCACCGCUCCUUAAGGAGUGGUGGCA<br>GGUAGGCGAAAACAUACCCCACACUCAACCCAUACCUAAAACCCCCA<br>CUCAACUCAAAACCAACCCAACAACAACAAACAAAAACUCACAACCAA<br>AUCUCACACACAACCAACCCCCAACCAACAAAACACAUAUAACACUCC<br>CUCACAAAUUCCCCCACACCCUAAAACAAAACCCCCCAACCAACAAU<br>CCUCAACCUACCAAAACAAACAACUACCCCAACAAAAACAAACACUAC<br>CCCCCAAUCUCUAUCCCCCACCCCCCAACACUUACCCCCCCUACUCC<br>CCAAAACCCCAUCCAAUACUCCCAACAAA | 0.0662 |

†Linguistic complexity(7,8) is defined as:

$$C = \prod_{k=1}^{w} U_k$$

where $U_k$ is the fraction of possible sequences observed for a *k*-mer and *w* is the maximum *k*-mer size. $U_k$ is:

$$U_k = \min[4^k, L - k + 1]$$

where *L* is the sequence length. *w* is adopted based on sequence length with $w$=3 for $L$<18, $w$=4 for 18≤$L$<67, $w$=5 for 67≤$L$<260, $w$=6 for 260≤$L$<1029, and $w$=7 for $L$≥1029.

**Supplementary Table 2. Primers (DNA oligo nucleotides) used for PCR to generate RNA-ID plasmids with various 5′UTR insert DNAs.**

| Primer name | Sequence (5' to 3') |
|---|---|
| 500-5UTR-200nt-F1 | TCAAATGTAATAAAACACCAAAAAACACACACACAACATAAAACAC AAAAACAAAACAACACCTAAACAACACACACACACACACACACA CACACACACACAAAACTACACAACAAACAACAACAAAACACAAAAC ACACACAACACACACACAACACAACAACAAAAAACACATACTAAAC ACAACACAAAAAACAC |
| 500-5UTR-200nt-F2 | CAAAAAACACATACTAAACACAACACAAAAAACACACAACACACAA CACACAACAACAAACCCCCACACCCCCCCCCTACAAAAACACAA AAAACACACACACTACACACATCATTACAACAATCACAACCTCACA CAACCCTATACCAAAACAAAAAACCCCTACACAAAAAACACACACA CAACTACACTAAATCTC |
| 500-5UTR-200nt-F3 | ACACAAAAAACACACACACAACTACACTAAATCTCACAAACAACAA CACAAAACACAACCTACACACACAAAACACAACCCCTCACTATTAT ACACACACAAAAAATACTAAACAAAAACAAAAAATAAAAAACACACA AAAACCCCCCTACTAAAACAAAAACAAAAATAAATAAAAAATACCCA AAATGTCTACTGAA |
| NR-5UTR-F1 | TCAAATGTAATAAAACACCAAAAAACACAC |
| 500NR-5UTR-F1 | ACACAAAAAACACACACACAACTAC |
| 500NR-5UTR-F2 | CAAAAAACACATACTAAACACAACAC |
| 500NR-5UTR-R1 | CTTCTAACTTCAGTAGACATTTTGGGTATTTTTTATTTATTTTTG |
| 500NR-5UTR-R2 | CGTTATTTCTTCTAACTTCAGTAGACATTTTGGG |
| 50NR-5UTR-F1 | CAAAACAACACCTAAAAATGTCTACTGAAGTTAGAAGAAATAACG |
| 50NR-5UTR-R1 | CTAACTTCAGTAGACATTTTTAGGTGTTGTTTTGTTTTTGTGTTTTAT G |
| 100NR-5UTR-F1 | ACACACACAAAACTACACAAAAAATGTCTACTGAAGTTAGAAGAAA TAAC |
| 100NR-5UTR-R1 | TAACTTCAGTAGACATTTTTTGTGTAGTTTTGTGTGTGTGTG |
| 150NR-5UTR-F1 | CACAACAAAAAATGTCTACTGAAGTTAGAAGAAATAACG |
| 150NR-5UTR-R1 | TTCAGTAGACATTTTTTGTTGTGTTGTGTGTGTGTTG |
| 150NR-5UTR-R2 | CGTTATTTCTTCTAACTTCAGTAGACATTTTTTGTTGTGTTG |
| 200NR-5UTR-F1 | CACACAACACAAAAATGTCTACTGAAGTTAGAAGAAATAAC |
| 200NR-5UTR-R1 | GTAGACATTTTTGTGTTGTGTGTTGTGTGTTTTTTG |
| 200NR-5UTR-R2 | GTTATTTCTTCTAACTTCAGTAGACATTTTTGTGTTGTGTG |
| 250NR-5UTR-F1 | AAAAACACAAAAAACACACAAAATGTCTACTGAAGTTAGAAGAAAT AAC |
| 250NR-5UTR-R1 | TAACTTCAGTAGACATTTTGTGTGTTTTTTGTGTTTTTGTAGG |
| 300NR-5UTR-F1 | CACAACCCTATACCAAAAATGTCTACTGAAGTTAGAAGAAATAAC |
| 300NR-5UTR-R1 | TAACTTCAGTAGACATTTTTGGTATAGGGTTGTGTGAGG |
| 350NR-5UTR-F1 | ACACAACTACACTAAATCTCAAAATGTCTACTGAAGTTAGAAGAAA TAAC |
| 350NR-5UTR-R1 | TTCTAACTTCAGTAGACATTTTGAGATTTAGTGTAGTTGTGTGTG |
| 400NR-5UTR-F1 | ACAACCCCTCAAAAATGTCTACTGAAGTTAGAAGAAATAAC |
| 400NR-5UTR-R1 | TTCAGTAGACATTTTTGAGGGGTTGTGTTTTGTG |
| 400NR-5UTR-R2 | CGTTATTTCTTCTAACTTCAGTAGACATTTTTGAGGGG |
| 450NR-5UTR-F1 | AAAAACAAAAAATAAAAAACAAAAATGTCTACTGAAGTTAGAAGAAA TAAC |
| 450NR-5UTR-R1 | TTCAGTAGACATTTTTGTTTTTTATTTTTTGTTTTTGTTTAGTATTTTT TG |
| SpeI-F1 | CCTATTTAAATGGTTGATGGATTACTAGTGG |
| SalI-R1 | GGACAGATTGTGTCGACAGGTAATGG |

| | |
|---|---|
| *URA3*-Replace-Ins-F1 | GGTTTATACATAATTTTACAACTCATTACGCACACTTAGTTTTGCTGGCCGCATCTTC |
| *URA3*-Replace-Vec-R1 | GAAGATGCGGCCAGCAAAACTAAGTGTGCGTAATGAGTTGTAAAATTATGTATAAACC |
| *URA3*-Replace-Vec-F1 | CACGTTCCTTATATGTAGCTTTCGACATTGTATGGATGGGGGTAATAGAATTGTATC |
| *URA3*-Replace-Ins-R1 | GATACAATTCTATTACCCCCATCCATACAATGTCGAAAGCTACATATAAGGAACGTG |
| StuI-remove-F1 | GTAACAAAGGAACCTAGAGGTCTTTTGATGTTAGCAG |
| StuI-remove-R1 | CTGCTAACATCAAAAGACCTCTAGGTTCCTTTGTTAC |
| 50SL2-300NR-5UTR-DNA1-R1 | AGCCCCCTCACCCAAGCAAGCTCGTAGAAATGAATGTTGTTGTTATAAATGAGGTGTTTG |
| 50SL2-300NR-5UTR-DNA1-R2 | TTTCGCATCTTAAGCCCCCTCACC |
| 50SL2-300NR-DNA2 -F1 | CGAAACTTAAGCCCTTTCGCCTAGGTGGGCAAAATCACAATACACAAAACCTACTAAACC |
| 50SL2-300NR-DNA2-F2 | ATGCGAAACTTAAGCCCTTTCGCC |
| RNAID-Vec-5'-350-R1 | TTGGGGTTTTAGGGAGGTTTTTATTGTTTTATTACATTTGAATAAGAAGTAATACAAACC |
| RNAID-Vec-5'-GAPDH-50 -F1 | AAAACCCCAATACTCCCTACTAATAAAACCCCCCGCTAGCCCCCCGGTTTCTATAAATTG |
| GAPDH-50-F2 | AAAACCCCAATACTCCCTACTAATAAAACCC |
| GAPDH-50RE-R2 | GGGTGTTTTATTACATTTGAATAAGAAGTAATACAAACCGAAAATG |
| GAPDH-50RE-F1 | ACAACCACACTCACATTTCCTACCCCCCCCATAAGCTAGCCCCCCGGTTTCTATAAATTG |
| GAPDH-50RE-F2 | TTATTCAAATGTAATAAAACACCCCCTAAATAATCACAACCACACTCACATTTCCTACCC |

**Supplementary Table 3. Primers used to generate gene deletion strain**

| Primer name | Sequence (5' to 3') | Amplicon used for |
|---|---|---|
| tif1-5UTR-F1 | GTAGAGTAAACTTCCACGCACATATTAGG | tif1 deletion |
| tif1-3UTR-R1 | CTCAACCATCCCAAACCTCGAG | tif1 deletion |
| tif2-5UTR-F1 | GCCTGATACCTGATGCCATCCC | tif2 deletion |
| tif2-3UTR-R1 | CAGCTAGTTACTCTTCACACACAAAG | tif2 deletion |
| tif3-5UTR-F1 | GATTAGTACCAGATTGAGCTCAGC | tif3 deletion |
| tif3-3UTR-R1 | TGCTGCAGTAACCTTCTTTGAAGG | tif3 deletion |
| tif4632-5UTR-F1 | CTGTCGCCAAGCTGTTCAAG | tif4632 deletion |
| tif4632-3UTR-R1 | CCAGAATACGTAAGCGTTCTGGATTC | tif4632 deletion |
| Tif-1-2-semiqPCR-F2 | CACCATCATGAAGGAATTCAGAAGTGG | tif1 or tif2 deletion confirmation |
| Tif-1-2-semiqPCR-R2 | GGATGGCAATTCTTCAATTTGAGTGGAG | tif1 or tif2 deletion confirmation |
| tif3-F1 | ATGGCTCCACCAAAGAAAACC | tif3 deletion confirmation |
| tif3-R2 | CTATTTCTTACCAACAACTTCCCAATTGTC | tif3 deletion confirmation |
| tif4632-F1 | CCACACCCGCAGCAAGCC | tif4632 deletion confirmation |

| | | |
|---|---|---|
| tif4632-R2 | TAATCACTGTCCCCATCGTTATTCATTAATG | tif4632 deletion confirmation |

**Supplementary Table 4. Primers used to generate mutant strains**

| No. | Primer name | Sequence (5' to 3') |
|---|---|---|
| 1 | Vec-DED1-site-F1 | CAAGCTTGTCGACGGAGCTCGTAATGCATCATTCTATACGTGTCATTCTG |
| 2 | Vec-DED1-site-R1 | CAGAATGACACGTATAGAATGATGCATTACGAGCTCCGTCGACAAGCTTG |
| 3 | DED1-TGA-3UTR-F1 | GTTCAAACAACTCTTCTTGGTGGTGATTTCAGACAAACTAGGGTGAGGATTC |
| 4 | DED1-TGA-3UTR-R1 | GAATCCTCACCCTAGTTTGTCTGAAATCACCACCAAGAAGAGTTGTTTGAAC |
| 5 | DED1-3UTR-KanMX-F1 | CCTGTATATTCGTTTTTGAATATACTTTGTTCCCAGCTTGCCTCGTCCCC |
| 6 | DED1-3UTR-KanMX-R1 | GGGGACGAGGCAAGCTGGGAACAAAGTATATTCAAAAACGAATATACAGG |
| 7 | KanMX500-vec-F1 | GGTGCGACAATCTATCGATTGTATGGCTAGCCATATGTATATCTCCTTCTTAAAG |
| 8 | KanMX500-vec-R1 | CTTTAAGAAGGAGATATACATATGGCTAGCCATACAATCGATAGATTGTCGCACC |
| 9 | Vec-KanMX-up-F1 | CAAGCTTGTCGACGGAGCTCAGCTTGCCTCGTCCCC |
| 10 | Vec-KanMX-up-R1 | GGGGACGAGGCAAGCTGAGCTCCGTCGACAAGCTTG |
| 11 | KanMX-3UTR500-F1 | CCGCCATCCAGTGTCGATTTCAGACAAACTAGGGTGAGGATTC |
| 12 | KanMX-3UTR500-R1 | GAATCCTCACCCTAGTTTGTCTGAAATCGACACTGGATGGCGG |
| 13 | 3UTR500-vec-F1 | GTATTCATTTCCTTCCTACATTTTCTCTCCGCTAGCCATATGTATATCTCCTTCTTAAAG |
| 14 | 3UTR500-vec-R1 | CTTTAAGAAGGAGATATACATATGGCTAGCGGAGAGAAAATGTAGGAAGGAAATGAATAC |
| 15 | Vec-TIF4631-site-F1 | CTTGGGCTGCAGGTCGACGACTCGATAACGACGTGAGAAACG |
| 16 | Vec-TIF4631-site-R1 | CGTTTCTCACGTCGTTATCGAGTCGTCGACCTGCAGCCCAAG |
| 17 | TIF4631-5UTR-ORF-F1 | CAAGGTAAGAGGACAACTGTAATTACCTATTACAATAATGACAGACGAAACTGCTCACCC |
| 18 | TIF4631-5UTR-ORF-R1 | GGGTGAGCAGTTTCGTCTGTCATTATTGTAATAGGTAATTACAGTTGTCCTCTTACCTTG |
| 19 | TIF4631-TAA-3UTR-F1 | GGCTCCTCCTCCAAAGGAAGAACCAGCTGCACCAACTTCTACCG |
| 20 | TIF4631-TAA-3UTR-R1 | CGGTAGAAGTTGGTGCAGCTGGTTCTTCCTTTGGAGGAGGAGCC |
| 21 | TIF4631-3UTR-HgrB500-F1 | CTTCGCCAGTCTTCGTTTACGGACATGGAGGCCCAG |
| 22 | TIF4631-3UTR-HgrB500-R1 | CTGGGCCTCCATGTCCGTAAACGAAGACTGGCGAAG |
| 23 | HgrB500-vec-F2 | GTGGATATGTCCTGCGGGTAAATGGATCCCCGGGCGAGCTCC |

| | | |
|---|---|---|
| 24 | HgrB500-vec-R2 | GGAGCTCGCCCGGGGATCCATTTACCCGCAGG ACATATCCAC |
| 25 | Vec-HgrB_salI-up-F1 | CTTGGGCTGCAGGTCGACGACATGGAGGCCCA G |
| 26 | Vec-HgrB_salI-up-R1 | CTGGGCCTCCATGTCGTCGACCTGCAGCCCAA G |
| 27 | HgrB-TIF4631-3UTR500-F1 | GAATGCTGGTCGCTATACTGCAGCTGCACCAAC TTCTACCG |
| 28 | HgrB-TIF4631-3UTR500-R1 | CGGTAGAAGTTGGTGCAGCTGCAGTATAGCGA CCAGCATTC |
| 29 | TIF4631-3UTR500-BamHI-vec-F1 | CAGAGGCCACGAAAGGAAAGGGGATCCCCGG GCGAGCTCC |
| 30 | TIF4631-3UTR500- BamHI-vec-R1 | GGAGCTCGCCCGGGGATCCCCTTTCCTTTCGT GGCCTCTG |
| 31 | Tif34Mut-Vec-ORF-F1 | GGGCTGCAGGTCGACTCTAGACAAATTATGGG ATGTGTCAAACGG |
| 32 | Tif34Mut-Vec-ORF-R1 | CCGTTTGACACATCCCATAATTTGTCTAGAGTC GACCTGCAGCCC |
| 33 | Tif34Mut-Mut-F2 | GGAATTCATTATTCTTGGTGGTGGTCGAGAGGC CAAGG |
| 34 | Tif34Mut-Mut-R2 | GGTGGTGACATCCTTGGCCTCTCGACCACC |
| 35 | Tif34Mut-3UTR1-HgrB500-F1 | CCCTTCGAGGTAATCTTCCGGTGGACATGGAG GCCCAG |
| 36 | Tif34Mut-3UTR1-HgrB500-R1 | CTGGGCCTCCATGTCCACCGGAAGATTACCTC GAAGGG |
| 37 | pSP64-XbaI-HgrB-F1 | GGGCTGCAGGTCGACTCTAGAGACATGGAGGC CCAG |
| 38 | pSP64-XbaI -HgrB-R1 | CTGGGCCTCCATGTCTCTAGAGTCGACCTGCA GCCC |
| 39 | HgrB-Tif34-3UTR500-F1 | GAATGCTGGTCGCTATACTGAGATCTTCCCTGT ATGCAGTTCC |
| 40 | HgrB-Tif34-3UTR500-R1 | GGAACTGCATACAGGGAAGATCTCAGTATAGCG ACCAGCATTC |
| 41 | Tif34-3UTR500-pSP64-BamHI-F1 | GAGGTTGTGTTATATAAGACAACGCTCGGATCC CCGGGCGAGCTCC |
| 42 | Tif34-3UTR500-pSP64-BamHI-R1 | GGAGCTCGCCCGGGGATCCGAGCGTTGTCTTA TATAACACAACCTC |
| 43 | Tif35Mut-Vec-ORF-F1 | GGGCTGCAGGTCGACTCTAGACATACAAGATTG AAGACGGTGTCAAG |
| 44 | Tif35Mut-Vec-ORF-R1 | CTTGACACCGTCTTCAATCTTGTATGTCTAGAGT CGACCTGCAGCCC |
| 45 | Tif35Mut-Mut1-F1 | CGTGATGATATGTGTACTTTGGCGATTATGCAA GTTAATG |
| 46 | Tif35Mut-Mut1-R1 | CATTTTCATTAACTTGCATAATCGCCAAAGTACA CATATC |
| 47 | Tif35Mut-Mut2-F1 | CAAGAGGTGCCGCCGCTGTTACCTTTTCGAG |
| 48 | Tif35Mut-Mut2-R1 | CTCGAAAAGGTAACAGCGGCGGCACCTCTTG |
| 49 | Tif35Mut-3UTR1-HgrB500-F1 | CAAAGCAAGGAAAAGCTCAGAGGACATGGAGG CCCAG |
| 50 | Tif35Mut-3UTR1-HgrB500-R1 | CTGGGCCTCCATGTCCTCTGAGCTTTTCCTTGC TTTG |
| 51 | HgrB-Tif35-3UTR500-F1 | GAATGCTGGTCGCTATACTGGCTGTCGTGAATA TGTCAAATACCG |

| | | |
|---|---|---|
| 52 | HgrB-Tif35-3UTR500-R1 | CGGTATTTGACATATTCACGACAGCCAGTATAG CGACCAGCATTC |
| 53 | Tif35-3UTR500-pSP64-BamHI-F1 | CTCGTTAAAGGAACTGTTGGTAGAATACCGGAT CCCCGGGCGAGCTCC |
| 54 | Tif35-3UTR500-pSP64-BamHI-R1 | GGAGCTCGCCCGGGGATCCGGTATTCTACCAA CAGTTCCTTTAACGAG |

**Supplementary Tables 5-11. Primers/template DNA combination for the PCR reaction that were used to produce fragment DNAs for Gibson assembly. Primer sequences are shown in Supplementary Table 3.**

**Table 5. Gibson assembly fragments used to generate pET24B-ded1-120-KanMX500.**

| DNA fragment to produce | Forward primer No. | Reverse primer No. | Template DNA |
|---|---|---|---|
| 5′UTR-ORF | 1 | 4 | gDNA from H5114-NSY5 |
| 3′UTR | 3 | 6 | gDNA from H5114-NSY5 |
| 5′end of hphMX | 5 | 8 | gDNA from H5120-NSY20 |
| Carrier vector | 7 | 2 | plasmid (pET24b) |

**Supplementary Table 6. Gibson assembly fragments used to generate pET24B-KanMX-ded1-120-3UTR500.**

| DNA fragment to produce | Forward primer No. | Reverse primer No. | Template DNA |
|---|---|---|---|
| hphMX | 9 | 12 | gDNA from H5120-NSY20 |
| 5′end of 3′UTR | 11 | 14 | gDNA from H5114-NSY5 |
| Carrier vector | 13 | 10 | plasmid (pET24b) |

**Supplementary Table 7. Gibson assembly fragments used to generate pSP64-tif4631-L614F-HgrB500.**

| DNA fragment to produce | Forward primer No. | Reverse primer No. | Template DNA |
|---|---|---|---|
| 5′UTR | 15 | 18 | gDNA from BY4741 |
| ORF | 17 | 20 | plasmid (B5608-pEP245) |
| 3′UTR | 19 | 22 | gDNA from BY4741 |
| 5′end of hphMX | 21 | 24 | gDNA from H5120-NSY20 |
| Carrier vector | 23 | 16 | plasmid (pSP64) |

**Supplementary Table 8. Gibson assembly fragments used to generate pSP64-tif4631-HgrB-3UTR500**.

| DNA fragment to produce | Forward primer No. | Reverse primer No. | Template DNA |
|---|---|---|---|
| hphMX | 25 | 28 | gDNA from H5120-NSY20 |
| 5′end of 3′UTR | 27 | 30 | gDNA from BY4741 |
| Carrier vector | 29 | 26 | plasmid (pSP64) |

**Supplementary Table 9. Gibson assembly fragments used to generate pSP64-tif34-Q258R-HgrB500.**

| DNA fragment to produce | Forward primer No. | Reverse primer No. | Template DNA |
|---|---|---|---|
| 5′UTR-ORF1 | 31 | 34 | gDNA from BY4741 |
| ORF2-3′UTR | 33 | 36 | gDNA from BY4741 |
| 5′end of HgrB | 35 | 24 | gDNA from H5120-NSY20 |
| Carrier vector | 23 | 32 | plasmid (pSP64) |

**Supplementary Table 10. Gibson assembly fragments used to generate pSP64-tif34-HgrB-3UTR500.**

| DNA fragment to produce | Forward primer No. | Reverse primer No. | Template DNA |
|---|---|---|---|
| hphMX | 37 | 40 | gDNA from H5120-NSY20 |
| 5′end of 3′UTR | 39 | 42 | gDNA from BY4741 |
| Carrier vector | 41 | 38 | plasmid (pSP64) |

**Supplementary Table 11. Gibson assembly fragments used to generate pSP64-tif35-KLF-HgrB500.**

| DNA fragment to produce | Forward primer No. | Reverse primer No. | Template DNA |
|---|---|---|---|
| ORF1 | 43 | 46 | gDNA from BY4741 |
| ORF2 | 45 | 48 | gDNA from BY4741 |
| ORF3-3′UTR | 47 | 50 | gDNA from BY4741 |
| 5′end of HgrB | 49 | 24 | gDNA from H5120-NSY20 |
| Carrier vector | 23 | 44 | plasmid (pSP64) |

**Supplementary Table 12. Gibson assembly fragments used to generate pSP64-tif35-HgrB-3UTR500.**

| DNA fragment to produce | Forward primer No. | Reverse primer No. | Template DNA |
|---|---|---|---|
| hphMX | 37 | 52 | gDNA from H5120-NSY20 |
| 5′end of 3′UTR | 51 | 54 | gDNA from BY4741 |
| Carrier vector | 53 | 38 | plasmid (pSP64) |

**Supplementary Table 13. Plasmids used for transformation to generate yeast mutant strains.**

| Mutant strain to generate | Plasmid | Restriction enzymes | Yeast strain to transform |
|---|---|---|---|
| ded1-120 | pET24B-ded1-120-KanMX500 and pET24B-KanMX-ded1-120-3UTR500 | SacI/NdeI | BY4741 |
| tif4632 deletion + tif4631-L614F | pSP64-tif4631-L614F-HgrB500 and pSP64-tif4631-HgrB-3UTR500 | SalI/BamH1 | Δtif4632 |
| tif34-Q258R | pSP64-tif34-Q258R-HgrB500 and pSP64-tif34-HgrB-3UTR500 | XbaI/BamH1 | BY4741 |
| tif35-KLF | pSP64-tif35-KLF-HgrB500 and pSP64-tif35-HgrB-3UTR500 | XbaI/BamH1 | BY4741 |

Plasmids were digested by restriction enzymes indicated and linear DNAs were agarose-gel purified and used for transformation.

**Supplementary Table 14. Primers used for RT-qPCR and 5′RACE**

| Primer name | Sequence (5' to 3') |
|---|---|
| tif1-2-semiqPCR-F1 | CGAAGAACCATCTGCCATTCAACAACG |
| tif1-2-qPCR-R1 | CCGGTCTTACCAGTACCAGATTGAGC |
| TFC-F1 | GCTGGCACTCATATCTTATCGTTTCACAATGG |
| TFC-R1 | GAACCTGCTGTCAATACCGCCTGGAG |
| TAF10-F1 | ATATTCCAGGATCAGGTCTTCCGTAGC |
| TAF10-R1 | GTAGTCTTCTCATTCTGTTGATGTTGTTGTTG |
| ALG9-F1 | CACGGATAGTGGCTTTGGTGAACAATTAC |
| ALG9-R1 | TATGATTATCTGGCAGCAGGAAAGAACTTGGG |
| GFP-qPCR-F4 | ATGGCCCTGTCCTTTTACCAGA |
| GFP-qPCR-R4 | AAGGACCATGTGGTCACG |
| RFP-qPCR-F5 | TAGCAGTTTGAGTACCTTCATATGG |
| RFP-qPCR-R5 | GTTCATATGGAAGGTTCAGTTAATGG |
| HIV-F1 | CTTCTGAAGATAAAGCAACAACAACAAG |
| 5RACE-R2 | CTCTCCACGGACAGAAAATTTGTG |
| NR-Lig-SacI-F2-2 | TTAACTATGAGCTCCTTCTGAAGATAAAGCAACAACAACAAG |
| NR-Lig-SacI-F3 | TTAACTATGAGCTCCTTCTGAAGATAAAG |
| M13R-Hiro | CACACAGGAAACAGCTATGACATG |
